# GD3 synthase deficiency disrupts Na^+^/K^+^-ATPase and plasma membrane Ca^2+^-ATPase function in mouse brain

**DOI:** 10.64898/2026.02.13.705793

**Authors:** Borna Puljko, Nikolina Maček Hrvat, Katarina Ilic, Ana Ujevic, Eva Josic, Mario Stojanović, Tadeja Režen, Klementina Fon Tacer, Damjana Rozman, Marta Balog, Marija Heffer, Svjetlana Kalanj-Bognar, Kristina Mlinac-Jerkovic

## Abstract

GD3 synthase (GD3S) is a key enzyme in the production of gangliosides, sialylated membrane glycosphingolipids with essential physiological roles in mammalian brains. To elucidate the molecular bases of neuropathological findings associated with GD3S deficiency, we performed a multilayered analysis focused on the functionality of ion transporters Na ^+^/K^+^-ATPase (NKA) and plasma membrane Ca^2+^-ATPase (PMCA) in the cortex and cerebellum of GD3S-deficient mice (GD3S^−/−^). We examined global transcriptomes, NKA and PMCA gene and protein expression, the influence of membrane lipid composition on lipid raft integrity, and the activity of both ATPases, pairing them with an exploratory principal component analysis. Transcriptomic data reveal that sets of genes involved in ion transport and membrane dynamics are differentially expressed in the absence of GD3S, whereas qRT-PCR data confirm changes in gene expression of specific NKA and PMCA subunits or isoforms. Altered protein expression and significantly lower activity of both NKA and PMCA were found in the cerebral cortex of GD3S^−/−^ mice. Detailed lipidomic analysis revealed segregation of cholesterol into lipid rafts, which may lead to disordered membrane lipid architecture in GD3S deficiency. Additionally, altered ganglioside composition was found to affect the activities of NKA and PMCA in the brain tissue of GD3S^−/−^ mice. Our results confirm that an imbalance in membrane ganglioside composition leads to significant alterations in ion transporter function. Experimental restoration of ATPase activity in cortical homogenates by administering exogenous b-series gangliosides may aid in developing therapeutic strategies targeting deficits in GD3S and other enzymes of ganglioside biosynthesis.

## 1. Introduction

Gangliosides are membrane sialoglycosphingolipids predominantly found in central nervous system (CNS) membranes and are involved in various processes, including signaling, differentiation, and synaptic function (Groux-Degroote et al., 2017; Mlinac-Jerkovic et al., 2026; Schnaar, 2019; Sipione et al., 2020; Yamaguchi and Kato, 2018; Yu et al., 2011). Because of their structural diversity and high concentration in the mammalian CNS compared to non-neural tissues, they are quantitatively significant components of the lipidome, glycome, and sialome of the brain and contribute to complex brain functions (Cohen and Varki, 2010). Gangliosides are classified into a, b-, and c-series, which display unique spatiotemporal expression and biophysical characteristics in brain tissue, especially regarding the assembly of lipid rafts (Groux-Degroote et al., 2017; Schnaar, 2016; Yu et al., 2011).

The sialyltransferase GD3 synthase (GD3S; alpha-N-acetylneuraminate alpha-2,8-sialyltransferase, EC 2.4.3.8), encoded by the *St8sia1* gene, is required for the biosynthesis of b- and c-series gangliosides by converting GM3 into GD3. GD3S has been more intensely studied in the context of tumorigenesis, where its overexpression is associated with metastasis and therapeutic resistance (Anand et al., 2025; Cao et al., 2023; Liu et al., 2018), however, its physiological function in the brain, where it is expressed at the highest level, as well as the consequences of its diminished expression, remains poorly understood. So far, only a handful of reports have described human conditions associated with ST8SIA1 mutations and the likely resulting reduction in GD3 synthase activity, although the findings remain conflicting (Husain et al., 2008; Ramagopalan et al., 2009).

GD3S-deficient mice (GD3S^−/−^), also referred to as *St8sia1* null mice, do not express the *St8sia1* gene. Whereas wild-type mice express four major brain gangliosides (GM1, GD1a, GD1b, and GT1b), the brains of *St8sia1* null mice lack GD1b and GT1b, instead expressing comparably increased levels of GM1 and GD1a. These mice display normal growth and overall morphology of nervous tissue (Wang et al., 2014); however, they exhibit signs of depressive behavior (Wang et al., 2014), impaired nerve regeneration after injury (Kittaka et al., 2008), memory impairments (Tang et al., 2021), auditory challenges (Orr et al., 2013), motor dysfunction, increased sensitivity in formalin testing (Ribeiro-Resende et al., 2014), thermal hyperalgesia and mechanical allodynia (Handa et al., 2005), as well as diminished visual acuity (Abreu et al., 2021). GD3S inactivation leads to physiological changes causing the observed phenotype, although the underlying molecular and cellular mechanisms remain only partially understood.

GD3S expression peaks during fetal development and early childhood, then declines with age (Svennerholm et al., 1989; Watanabe et al., 1996; Yamamoto et al., 1996). It often increases in malignant conditions, where rapidly growing tumor cells exhibit high levels of GD3 ganglioside. Studies have shown that inactivating GD3S can reduce tumor formation and metastasis, making it a potential therapeutic target (Cao et al., 2023; Kasprowicz et al., 2022; Liu et al., 2018; Ohkawa et al., 2021; Zhang et al., 2021). Additionally, inhibition of GD3 synthase has been proposed as a therapeutic approach for neurodegeneration, as amyloid-like neuropathology was found to be diminished in GD3S^−/−^ mice (Bernardo et al., 2009). Therefore, understanding the physiological effects of GD3S inactivation on cells and tissues is crucial, given the unclear consequences of complete GD3S loss.

In this study, we examined two differently organized types of cortical tissue - neocortex and cerebellum - of GD3S^−/−^ mice at the transcriptome, proteome, lipidome and enzyme activity levels in order to elucidate the molecular mechanisms underlying neuropathology due to GD3S depletion. We previously showed that GD3S deficiency affects both cholesterogenic gene expression (Mlinac et al., 2012) and fatty acid metabolism (Puljko et al., 2024). Furthermore, using other models, our group has previously reported on the importance of ganglioside composition for the proper function of ion transporters Na^+^/K^+^-ATPase (NKA, EC 7.2.2.13) and plasma membrane Ca^2+^-ATPase (PMCA, EC 7.2.2.10) (Ilic et al., 2021; Puljko et al., 2021). Based on the whole-genome expression results presented here and our previous findings, we focused on analyzing membrane lipid distribution in relation to NKA and PMCA, both implicated in several neurological disorders. NKA is a ubiquitous ATP-hydrolyzing pump, responsible for upholding the electrochemical gradient (Benarroch, 2011; Kaplan, 2002; Pirahanchi et al., 2021). Disturbances in ion homeostasis affect neuronal excitability and synaptic transmission, and changes in NKA activity have been reported in various neurological disorders (Bøttger et al., 2012, 2011; Pietrobon and Conti, 2024; Sun et al., 2022; Ygberg et al., 2021). PMCA is responsible for transporting calcium ions across the plasma membrane, which is essential for regulating intracellular calcium levels and modulating diverse cellular processes, including signaling and synaptic transmission. PMCA exists in four main isoforms, PMCA1-4, which are the result of alternative splicing. These isoforms have distinct structures, tissue-specific expression patterns, and kinetic parameters, enabling functional specialization across diverse cell types and distinct catalytic properties (Boczek et al., 2019; Domi et al., 2007; Kip and Strehler, 2003; Krebs, 2015). PMCA dysfunction has also been linked to several neurological diseases (Berrocal and Mata, 2023; Gu et al., 2025).

In this study, we demonstrate that the altered ganglioside composition resulting from GD3S deficiency causes significant perturbations in plasma membrane organization, which in turn impair the function of the ion transporters NKA and PMCA. Furthermore, we show that the compromised catalytic activity of these ATPases can be restored by exogenous treatment with the gangliosides GD1b and GT1b, which are absent in the GD3S^−/−^ mouse model.

## 2. Materials and methods

### 2.1. Experimental animals

A total of 38 male mice (5-7 months old) were used: 16 GD3S^−/−^ (*St8sia1 null*) mice and 22 wild-type (WT) controls (B6;129S, the same genetic background as the null animals). Animals were bred and maintained under certified conditions (HR-POK-005; 12/12 h light/dark cycle, constant temperature and humidity, ad libitum food and water, enriched environment). All procedures were approved by the institutional and national authorities (Ethics Committee of the University of Zagreb for the Croatian Science Foundation projects HRZZ-IP-2016-06-8636 and HRZZ-IP-2014-09-2324, Ethics Committee of the Faculty of Medicine Osijek and Croatian Ministry of Agriculture under class number 641-01/15-01/49, reference number 525-61-07-15-07) and complied with appropriate animal welfare regulations (Animal Protection Act 102/2017; Ordinance 55/2013) and EU Directive 2010/63/EU, including ARRIVE 2.0 guidelines and the 3R principles. The animals were euthanized by cervical dislocation, brains rapidly removed, cortex and cerebellum dissected, frozen in liquid nitrogen, and stored at −80⍰°C. At least three biological replicates were used for every experiment. A schematic overview of the study is shown in Supplementary Figure 1.

### 2.2. Microarray analysis

Whole-genome expression was assessed in cerebellar tissue from GD3S^−/−^ mice compared to WT mice. The tissue was homogenized in TRI Reagent (Molecular Research Center, #RT111), and total RNA was extracted according to the manufacturer’s guidelines. The concentration and quality of the extracted RNA were determined spectrophotometrically and with the RNA 6000 Nano Kit on a Bioanalyzer 2100 (Agilent Technologies, Palo Alto, CA, USA). Gene expression analysis was performed using 1 x 22K Whole Mouse Genome Oligo Microarrays (Agilent Technologies, #G4122A) according to the manufacturer’s instructions. A two-color indirect experimental design was employed, where each mouse sample (GD3S^−/−^ or WT) was hybridized with a reference sample created by mixing equal amounts from all mouse samples. One microgram of total RNA from six mouse samples and six reference samples was used for cDNA synthesis, followed by cRNA synthesis with cyanine-3 and -5 labeled CTP (Perkin Elmer/NEN Life Sciences, #NEL580001EA and #NEL581001EA) using the Low RNA Input Linear Amplification Kit Plus, Two-Color (Agilent Technologies, #5188-5339). Labeled cRNA samples were purified using the RNeasy Mini Kit (Qiagen GmbH, #74004). Each mouse sample was combined with a reference sample to prepare the hybridization mix using the Gene Expression Hybridization Kit (Agilent Technologies, #5188-5242) according to the manufacturer’s guidelines. Samples were hybridized to microarrays in hybridization chambers in an Agilent oven at 65°C for 17 hours. After hybridization, the microarrays were washed with the Expression Wash Buffer Kit (Agilent Technologies, #5188-5327) according to the manufacturer’s instructions. The microarrays were scanned using the LS 200 scanner (Tecan Group Ltd.), and images were analyzed with GenePixPro 6 (Molecular Devices LLC). The microarray data have been deposited in the GEO database under accession GSE318770. Lowess normalization and classification of differentially expressed genes (DEGs) were conducted using BRB-ArrayTools Version 3.7.0 beta_2, developed by Dr. Richard Simon and Amy Peng Lam. Class comparisons were performed between groups using a significance threshold of α = 0.001 and an FDR < 0.165, excluding one sample due to its low overall signal. Enrichment analysis of transcription factors among DEGs was performed using Enricher (Chen et al., 2013; Kuleshov et al., 2016; Xie et al., 2021) with the ENCODE and ChEA databases, and TRANSFAC (geneXplain GmbH, Germany) was used to analyze the promoters of DEGs.

### 2.3. Reverse transcription quantitative PCR (RT-qPCR)

Total RNA was extracted from cortical and cerebellar brain tissue samples using the GeneJET RNA Purification kit (Thermo Fisher Scientific, #K0702). Reverse transcription was performed with the High-Capacity cDNA Reverse Transcription Kit alongside RNase Inhibitor (Applied Biosystems, #4388950) on an equal quantity of total RNA, followed by DNase I treatment (Sigma-Aldrich, #EN0521). For RT-qPCR analysis, 2 ng of each sample’s cDNA was analyzed using the Applied Biosystems™ SYBR™ Green Universal Master Mix (Applied Biosystems, #4309155). The primer sequences are listed in Supplementary Table 1. Primers were designed using the NCBI Primer-BLAST Software (Ye et al., 2012) and validated for specificity and efficiency (Bookout et al., 2006). Real-time fluorescence detection was performed using a 7900 HT Real-Time PCR System (Applied Biosystems), and data were quantified using the relative quantification method (ΔΔCt), normalized to the housekeeping gene *Actb*, and presented to compare relative target gene expression between groups (Bookout et al., 2006).

### 2.4. Western blotting

Cortical and cerebellar brain tissue samples were thawed on ice and homogenized in ddH2O to a 10% (v/w) homogenate. The total protein concentration was measured using the Pierce™ BCA Protein Assay Kit (Thermo Fisher Scientific #23225) following the manufacturer’s instructions. Proteins were separated on SDS-PAGE (15 µg of each sample per lane) using the TGX Stain-Free™ FastCast™ 12% Acrylamide Kit (Bio-Rad #1610185). On the same gel, WT and GD3S^−/−^samples were run in parallel. Following electrophoresis, gels were activated by UV light for 5 minutes using a ChemiDoc MP Imaging System (Bio-Rad). Proteins were transferred to PVDF membranes using the Mini-Blot Module (Invitrogen #B1000), and the membranes were imaged with the ChemiDoc MP Imaging System. The membranes were blocked in phosphate-buffered saline containing 0.1% Tween-20 (PBST) with 5% skim milk for 1 hour at room temperature, then incubated overnight at 4 °C with primary antibodies diluted in the same blocking solution. The following primary antibodies were used at the indicated dilutions: anti-panNKA (Abcam #ab300507, RRID: AB_3713501; 1:2500); anti-α 1 NKA (Abcam #ab76020, RRID: AB_1310695; 1:100,000); anti-panPMCA (Merck #MABN1802, RRID: AB_3713503; 1:2,000); anti-PMCA1 (Abcam #ab190355, RRID: AB_2893200; 1:1,000); anti-PMCA2 (Abcam #ab3529, RRID: AB_303878; 1:1,000); anti-PMCA3 (Novus Biologicals #NBP1-59465, RRID: AB_11016874; 1:1,000); anti-PMCA4 (Abcam #ab2783, RRID: AB_303296; 1:1,000); anti-transferrin receptor (TfR; Thermo Fisher Scientific #13-6800, RRID: AB_2533029; 1:1,000); anti-Flotillin1 (BD Biosciences #610821, RRID: AB_398140; 1:1,000); and cholera toxin subunit B (CTB which recognizes the ganglioside GM1; Thermo Fisher Scientific #ab2783, RRID: AB_10971380; 1:50,000). After washing in PBST, blots were exposed to peroxidase-labelled secondary antibodies (Jackson ImmunoResearch #715-035-150; RRID: AB_2340770, #711-035-152; RRID: AB_10015282, #713-035-147; RRID: AB_2340710, #705-035-003; RRID: AB_2340390) diluted 1:50,000 in PBST. Bands were visualized using the SuperSignal™ West Femto Maximum Sensitivity Substrate (Thermo Fisher Scientific, #34095) and imaged with the ChemiDoc MP Imaging System (Bio-Rad). Immunoreactivity was normalized to total protein per lane using *stain-free* imaging (Maloy et al., 2022; Rivero-Gutiérrez et al., 2014), and band intensities were quantified with ImageLab 6.6.1 software (Bio-Rad).

### 2.5. Lipid rafts isolation

Lipid rafts (LR) and bulk membrane fractions (non-LR; nLR) were isolated following established protocols (Mlinac-Jerkovic et al., 2021). Frozen samples of cortical and cerebellar brain tissue were thawed on ice. Approximately 70 ± 5 mg of tissue was homogenized in a detergent-free homogenization buffer (50 mM Tris (pH 8), 150 mM NaCl, 1 mM MgCl_2_, 1 mM CaCl_2_, and protease inhibitors: 1 mM phenylmethylsulfonyl fluoride (PMSF), 5 mM NaF, 1 mM Na_3_VO_4_, and 1% protease inhibitor cocktail) containing 0.32 M sucrose, using 30 strokes in a Potter-Elvehjem glass homogenizer with a Teflon pestle. The homogenate was centrifuged at 1,000 g for 15 minutes at +4°C, and the postnuclear supernatant was ultracentrifuged at 100,000 g for 30 minutes at +4°C (using a Beckman Optima XL-80 K ultracentrifuge; 50.4 Ti rotor). The resulting pellet was homogenized in a buffer containing 1% Brij O20, transferred to a pre-cooled SW 28.1 Ti ultracentrifuge tube, and gently mixed with 600 µL of 85% sucrose (w/v) prepared in the same homogenization buffer with 1% Brij O20. This mixture was carefully overlaid with 10 mL of 35% sucrose (w/v) in homogenization buffer containing 1% Brij 20 and 4 mL of 3% sucrose (w/v) in the same buffer. The tubes were ultracentrifuged at 141,000 x g for 18 hours at +4°C (Beckman Optima XL-80 K; SW 28 rotor). A distinct band of lipid rafts was observed at the interface between the 3% and 35% sucrose layers after centrifugation. Ten fractions were then collected from top to bottom and stored at −80°C. The successful isolation of rafts was confirmed by Western blotting using established LR and nLR markers (Persaud-Sawin et al., 2009) (Supplementary Figure 2).

### 2.6. Cholesterol assay

Cholesterol levels in tissue homogenates from the cerebral cortex and cerebellum, as well as in LR and nLR fractions from both brain regions, were assessed using the Amplex Red Cholesterol Assay Kit (Thermo Fisher Scientific, #A12216) according to the manufacturer’s guidelines and detected using the GloMax Discoverer plate reader (Promega).

### 2.7. ATPase activity assays

Na^+^/K^+^-ATPase and plasma membrane Ca^2+^-ATPase activities were measured in cortical and cerebellar homogenates by quantifying inorganic phosphate (Pi) released from ATP hydrolysis, following established protocols (Puljko et al., 2021). Three experimental setups were used: (i) endogenous enzyme activity was measured in cortical and cerebellar homogenates from WT and GD3S^−/−^ mice; (ii) potential modulatory effects of exogenous gangliosides (GM1, GD1a, GD1b, and GT1b; Sigma-Aldrich) were evaluated in WT cortical homogenates using a time- and concentration-dependent screening assay (varying preincubation for 15, 30, 60, or 120 minutes at concentrations of 10^−2^, 10^−3^, 10^−4^, 10^−6^, 10^−7^ and 10^−8^⍰mM); and (iii) GD3S^−/−^ cortical homogenates were tested after 2-hour preincubation with GD1b or GT1b at concentrations determined from the preceding screening. 10% water homogenates were prepared by disrupting the tissues with 15 strokes using a Potter-Elvehjem glass homogenizer with a Teflon pestle. All steps were carried out on ice unless stated otherwise. Reaction mixtures contained 50⍰µg of total protein in a buffer composed of 30⍰mM Tris-HCl (pH 7.4), 3⍰mM MgCl_2_, 100⍰mM NaCl, 20⍰mM KCl, and 3⍰mM ATP. Tested ganglioside concentrations were 10^−2^, 10^−3^, 10^−4^, and 10^−6^⍰mM. Samples were preincubated at 37⍰°C for 10 minutes prior to ATP addition. Reactions were allowed to proceed for 15 minutes, then stopped by the addition of 10% (w/v) trichloroacetic acid. After centrifugation (20,000 × g, 10 minutes, 4 °C), the supernatants were incubated with freshly prepared chromogenic reagent containing 150 mM ferrous sulfate in 1% (w/v) ammonium heptamolybdate dissolved in 0.6 M H_2_SO_4_, for 45 minutes. The Pi concentration was determined by measuring absorbance at 700 nm using a Varian Cary 100 Bio spectrophotometer (SpectraLab Scientific Inc.). Specific NKA and PMCA activities were calculated as the difference between total and ouabain-sensitive or carboxyeosin-sensitive hydrolysis, respectively, and expressed as micromoles of Pi per hour per milligram of protein (µmol(Pi)/(mg (protein)·h)).

### 2.8. Ganglioside extraction and dot blotting

Gangliosides were isolated from cortical tissue using previously established protocols (Mlinac et al., 2013). Tissue was homogenized in cold distilled water using a glass-Teflon Potter-Elvehjem homogenizer. Lipids were then extracted with a chloroform: methanol (1:2, v/v) mixture. The solvent ratio was adjusted to 1:1:0.7 (chloroform:methanol:water) to facilitate phase separation. The upper layer, containing gangliosides, was collected and further purified by Sephadex G-25 gel filtration. For quantification, 1⍰µL of ganglioside fractions was spotted onto nitrocellulose membranes for dot blot analysis. Membranes were blocked with 5% non-fat powdered milk in PBST for 1 hour at ambient temperature, then incubated overnight at 4°C with primary antibodies: CTB to detect GM1 (Thermo Fisher Scientific #ab2783, RRID: AB_10971380; 1:50,000) and anti-GD1a (0.32 μg/mL, provided by Dr. Ronald L. Schnaar, Johns Hopkins University). After washing, membranes were exposed to HRP-conjugated secondary antibodies (Jackson ImmunoResearch #715-035-150; RRID: AB_2340770; 1:50,000) for 1 hour at ambient temperature. Detection was performed via chemiluminescence, using the same procedure described for Western blotting, Section 2.4.

### 2.9. Statistical analysis and study design

All statistical analyses were performed using GraphPad Prism version 10.2.0 (GraphPad Software) and Microsoft Excel. Data are reported as mean ± standard error of mean (SEM). Normality was assessed with the Shapiro-Wilk test. For data following a normal distribution, an unpaired two-tailed t-test was used; otherwise, the Mann-Whitney test was applied. Statistical significance was set at p < 0.05. Sample sizes were estimated a priori with G*Power based on effect sizes from preliminary transcriptomic and enzyme activity data (α = 0.05; Cohen’s d: PMCA = 3.64, power = 0.97; NKA = 3.10, power = 0.94). Five biological replicates per genotype were used for RT-qPCR, proteomic, and biochemical analyses, enabling integrated molecular profiling across matched samples. All assays were performed in technical duplicates or triplicates, depending on the sample availability. Relationships among transcripts, proteins, and enzyme activities were assessed using Pearson or Spearman’s rank correlation on z-score-standardized data. Exploratory principal component analysis (PCA) and principal component regression (PCR) were performed on a subset of animals using centered z-score standardized membrane-associated measurements (GraphPad Prism version 10.2.0). PCA was used for dimensionality reduction and visualization of multivariate structure; PCR used enzyme activity as the outcome variable and principal components as predictors. Biplots were generated with 95% confidence ellipses and loading vectors to illustrate group separation and variable contributions. In line with best practice, no *post hoc* analysis was done.

## 3. Results

### 3.1. Microarray analysis revealed differentially expressed genes in GD3S^−/−^ mice

To gain initial insights into how altered ganglioside composition affects molecular pathways in GD3S^−/−^ mice, we performed genome-wide microarray analysis of the cerebellar tissue The heatmap in Figure 1A highlights the top 25 differentially expressed genes (DEGs) between GD3S^−/−^ and wild-type (WT) mice, filtered for statistical significance (*p < 0.001) and a threshold of |log_2_(fold change) = 0.5|, generated using ShinyGO v0.82 (Ge et al., 2020). Among the DEGs, we observed changes in genes linked to ion transport, membrane dynamics, and cellular metabolism. Notably, *Atp1b3*, encoding the β3 regulatory subunit of NKA, and *Atp2b2*, encoding the PMCA2 isoform, showed different expression in the cerebellum of GD3S^−/−^ mice, suggesting possible alterations in membrane pump composition and activity. Supplementary Table 2 presents additional data on DEGs involved in neuronal signaling, membrane transport, RNA processing, and cellular architecture. To further explore potential regulatory mechanisms related to the observed transcriptional changes, we performed transcription factor enrichment analysis on the DEGs and gene ontology (GO) enrichment analysis. The GO enrichment analysis (Figure 1B) highlights overrepresented processes, primarily related to ion transport and gene transcription (e.g., RNA binding and nucleic acid binding). To illustrate potential interactions between DEGs and enriched transcription factors, and to provide insight into broader transcriptional reprogramming in GD3S deficiency, we performed network visualization in Cytoscape (Supplementary Figure 3) using the STRING database for DEGs and selected significantly enriched transcription factors using Enricher (Chen et al., 2013). Microarray findings were further validated by qRT-PCR analyses of *Kras* gene expression in cerebellar tissue (Supplementary Figure 4), as well as RT-qPCR for all NKA subunits and PMCA isoforms, which are discussed in subsequent sections. Following RT-qPCR analyses, this investigation was extended to cortical tissue, enabling a comparison of regional expression patterns in brain regions with distinct ganglioside expression profiles (Lee et al., 2024). Since transcriptome analysis results accentuated processes related to ion transport (Figure 1B) and considering previous research on the effect of gangliosides on ion transport (Ilic et al., 2021; Jiang et al., 2014; Leon et al., 1981; Puljko et al., 2021) as well as recent reports on the ganglioside interactome in human cells (Zhang et al., 2024), we proceeded with the in-depth analysis of two plasma membrane ion transporters, NKA and PMCA.

**Figure 1.**
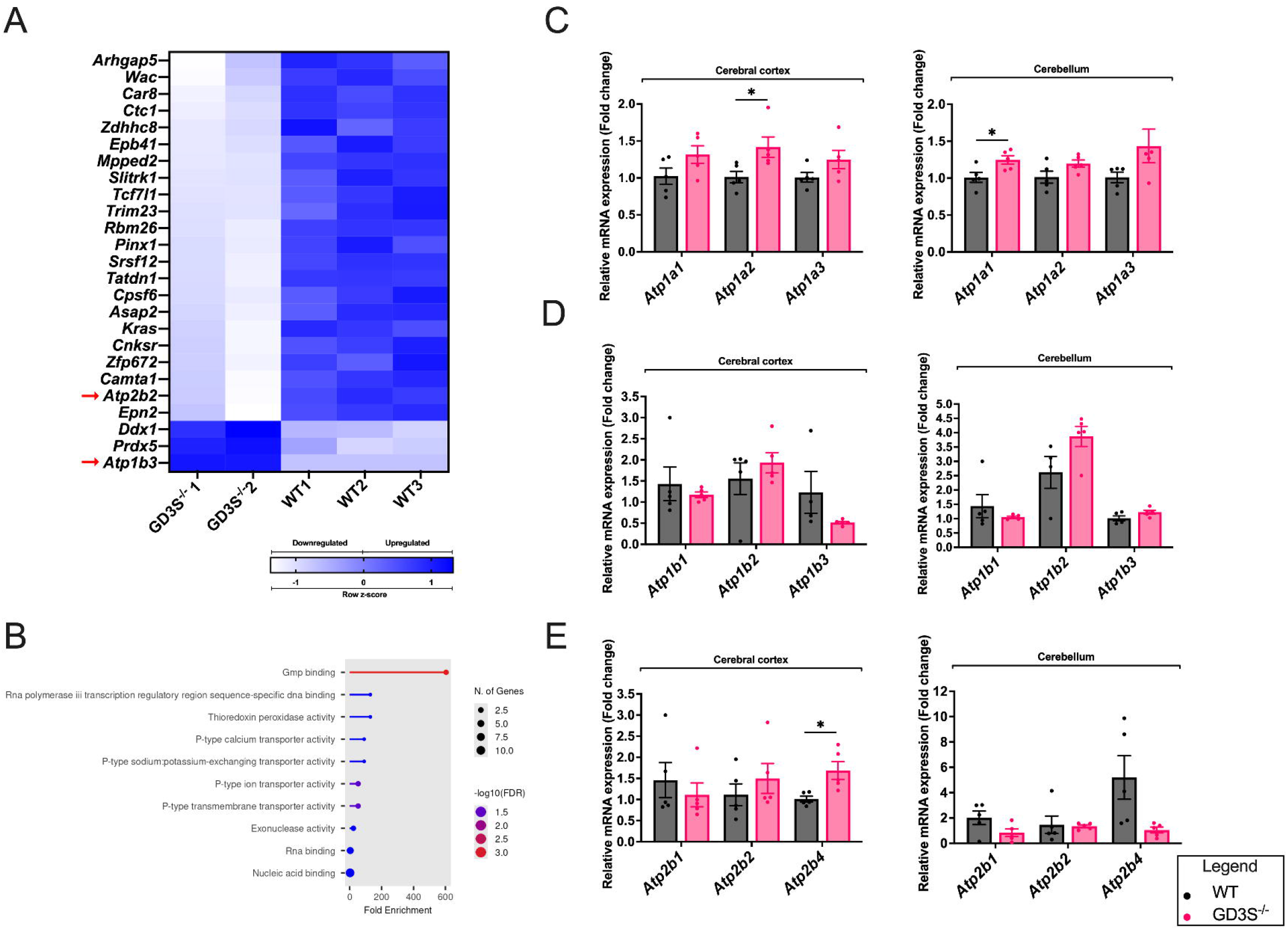
Differential gene expression and qRT-PCR validation in the cerebral cortex and cerebellum of GD3S^−/−^ mice. (A) Heatmap of top 25 DEGs with the lowest FDR values in the cerebellum of GD3S^−/−^ vs. WT mice, filtered for p < 0.001 and |log_2_(fold change)| ≥ 0.5; n = 2 GD3S^−/−^, 3 WT. Genes selected for further analysis in this study are marked with red arrows. (B) GO enrichment analysis of 25 DEGs in GD3S^−/−^ mice cerebella, highlighting overrepresented processes related to ion transport. Circle size represents gene count; color indicates fold enrichment. Analysis performed using ShinyGO v0.82. (C, D) qRT-PCR analysis of (C) NKA catalytic subunits (*Atp1a1, Atp1a2, Atp1a3*) and (D) regulatory β subunits (*Atp1b1, Atp1b2, Atp1b3*) in the cerebral cortex and cerebellum of GD3S^−/−^ and WT mice. Bars represent mean ± SEM from n = 5 mice per group; * = p < 0.05, two-tailed unpaired *t*-test. (E) qRT-PCR analysis of genes encoding PMCA isoforms (*Atp2b1, Atp2b2, Atp2b4*) in the cerebral cortex and cerebellum of GD3S^−/−^ and WT mice. Bars represent mean ± SEM from n = 5 mice per group; * = p < 0.05, two-tailed unpaired *t*-test.

### 3.2. GD3S deletion affects expression of ATPase-related genes

We examined the expression of 10 genes involved in ion transport in the cortex and cerebellum of GD3S^−/−^ and WT mice. *Atp1a1, Atp1a2*, and *Atp1a3* (Figure 1C), which encode the catalytic α-subunits of Na+/K+-ATPase (NKA), exhibited higher expression in the cortex of GD3S^−/−^ mice compared to WT mice. Specifically, *Atp1a2* levels were significantly higher in GD3S^−/−^ mice (p < 0.05, two-tailed unpaired t-test). A similar pattern of increased gene expression was observed in the cerebellum, with a significant difference in *Atp1a1* expression. There was no significant regional change in the expression of *Atp1b1, Atp1b2*, and *Atp1b3*, which encode the β-subunits of NKA (Figure 1D). *Atp2b1, Atp2b2*, and *Atp2b4* (Figure 1E) encode different plasma membrane Ca^2+^-ATPase (PMCA) isoforms and the levels of the latter were significantly higher in GD3S^−/−^ mice cortices (p < 0.05). However, in the cerebellum of GD3S^−/−^ animals, the expression of all three isoforms was decreased (p < 0.05) (Figure 1E). The expression of *Atp2b3* was bellow level of detection, because amplification of *Atp2b3* transcripts in both brain regions required more than 45 cycles, in both brain regions. Together, these results demonstrate not only the differences in expression between WT and GD3S^−/−^ mice but also highlight region-specific differences in ion transporter gene expression in control animals.

### 3.3. Region-specific alterations in NKA and PMCA mRNA lead to similar patterns in protein levels in GD3S^−/−^ mice

To assess whether mRNA changes resulted in altered protein levels, we conducted western blot analyses of NKA and PMCA isoforms in the cerebral cortex and cerebellum of GD3S^−/−^ and WT mice (Figure 2A-D). In the cortex, total NKA levels detected with a panNKA antibody were significantly higher in GD3S^−/−^ mice compared to WT (p < 0.001, Student’s t-test), aligning with increased transcript levels of all α subunits found by qRT-PCR (Figure 1C). However, the gene expression of the α1 NKA subunit was significantly lower in the GD3S^−/−^ cortex (p < 0.05). In cerebellar tissue, α1 NKA also showed a decreased expression trend (Figure 2C). Total PMCA protein levels (panPMCA) did not differ significantly between genotypes in either brain region investigated; however, isoform-specific changes were observed (Figure 2D). PMCA1 was significantly reduced in GD3S^−/−^ cortex (p < 0.001), while PMCA3 expression was considerably decreased in both regions (p < 0.001). PMCA2 remained unchanged across regions. Overall, the Western blot results indicate that GD3S deficiency induces region-specific changes in ion transporter protein levels that partly reflect transcriptomic alterations.

**Figure 2.**
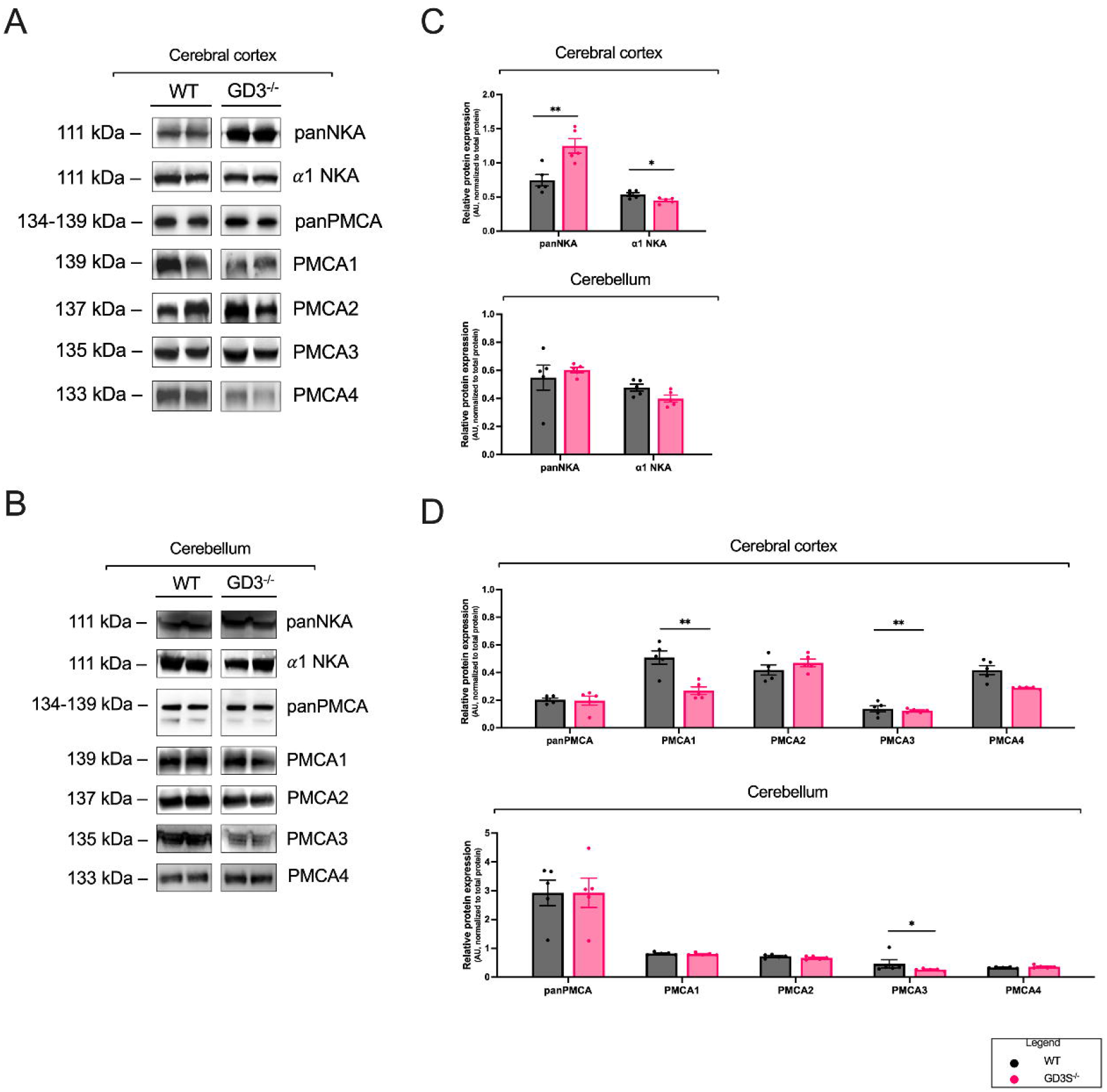
Regional protein expression of Na+/K+-ATPase (NKA) subunits and PMCA isoforms in the cerebral cortex and cerebellum of WT and GD3S^−/−^ mice. (A, B) Representative Western blots demonstrate the expression of pan-NKA, α1-NKA, pan-PMCA, and PMCA1-4 in cerebrocortical (A) and cerebellar (B) homogenates from WT and GD3S^−/−^ mice. Two samples are presented per group for clarity. (C) Quantification of pan-NKA and α1-NKA expression reveals significant differences between WT and GD3S^−/−^ mice in the cerebral cortex and cerebellum. (D) Quantification of pan-PMCA and PMCA1-4 isoforms demonstrates significant differences between WT and GD3S^−/−^ mice in the cerebral cortex and cerebellum. All samples were processed on the same gel and analyzed based on total protein normalization per well (*stain-free* imaging). Bars represent mean ± SEM, and n = 5 animals per group. Statistical comparisons were performed using two-tailed unpaired *t*-tests or Mann-Whitney tests as appropriate; *p < 0.05, **p < 0.001. AU, arbitrary units.

### 3.4. GD3S deficiency causes membrane redistribution of NKA and PMCA isoforms

To examine how GD3S deficiency affects lateral membrane organization and the distribution of NKA and PMCA, we quantified the distribution of ATPases and cholesterol in lipid raft (LR) and non-lipid raft (nLR) membrane domains in the cerebral cortex and cerebellum. Western blot immunoreactivity from LR and nLR fractions was used to determine raft partitioning coefficients (K_p, raft_), which are defined as the ratio of immunoreactivity in the LR fraction to that in the nLR fraction (I_LR_/I_nLR_), normalized to wild-type (WT) K_p, raft_ values (Lorent et al., 2017). Values of K_p, raft_ > 1 indicate a preference for lipid raft localization, while values < 1 suggest non-raft distribution. In the cortex, GD3S^−/−^ samples showed reduced raft preference for most proteins, with notable decreases in PMCA1 (0.52), PMCA2 (0.43), PMCA3 (0.37), and α1NKA (0.42). Conversely, the raft enrichment of PMCA4 (2.57) increased (Figure 3A). In contrast, the cerebellum showed a partial reversal of this trend; GD3S^−/−^ mice-derived fractions had increased raft association of α1NKA (1.17); however, most other proteins, including all PMCA isoforms, remained below a K_p, raft_ value of 1 (Figure 3B). Total cholesterol quantification in whole-tissue homogenates revealed only slightly elevated levels, not statistically significant, in both the cortex and cerebellum of GD3S^−/−^ mice (Figures 3C). However, domain-specific analysis showed significantly increased cholesterol in LR fractions from GD3S^−/−^ cortex and cerebellum, while nLR cholesterol levels were similar to the WT levels (Figure 3D). Furthermore, dot blots detecting a-series gangliosides GM1 and GD1a (Figure 3E) confirm their higher levels in cortical and cerebellar tissue of GD3S^−/−^ mice. Overall, these findings suggest that GD3S deficiency rearranges membrane composition by modulating both protein partitioning and lipid content, potentially affecting ATPase activity through changes in membrane microdomain organization and distribution.

**Figure 3.**
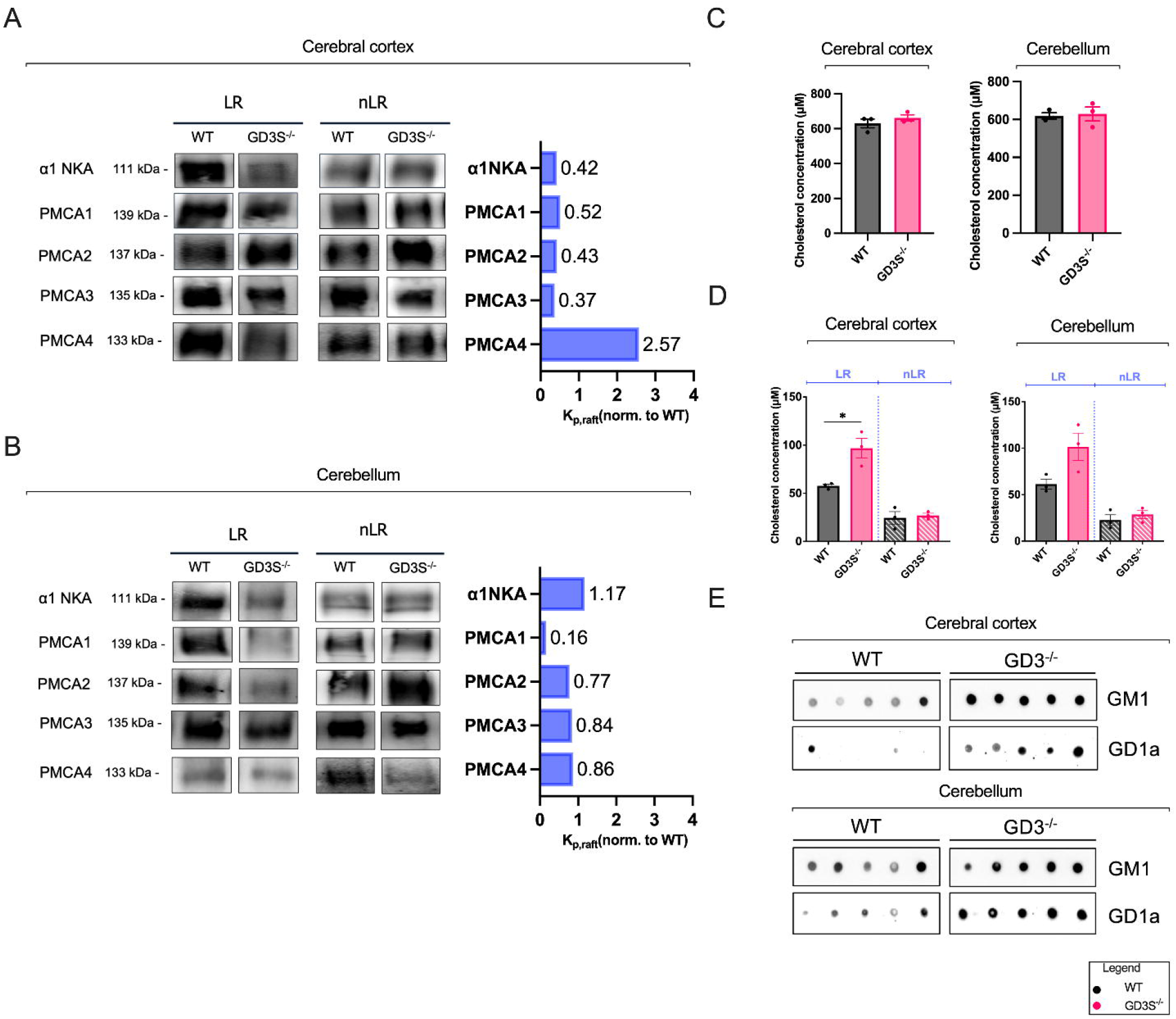
Membrane subdomain distribution patterns of NKA and PMCA isoforms, and membrane lipid composition in cerebral cortex and cerebellum of WT and GD3S^−/−^ mice. (A, B) Western blots showing α1-NKA and PMCA isoforms 1-4 in lipid raft (LR) and non-raft (nLR) fractions from cerebral cortex (A) and cerebellum (B) of WT and GD3S^−/−^ mice. Corresponding bar graphs to the right show K_p, raft_ values normalized to WT for each protein (n = 3 per group). Values > 1 indicate raft-preferencing localization; values <1 indicate non-raft preference. C) Quantification of cholesterol concentration (µM) in whole homogenates from cerebral cortex and cerebellum of WT and GD3S^−/−^ mice (n = 3 per group). (D) Cholesterol content measured in LR and nLR fractions from cortex and cerebellum of WT and GD3S^−/−^ mice (n = 3 per group). All bars represent mean ± SEM. Statistical comparisons were performed using a two-tailed unpaired *t*-test (*p < 0.05). (E) Dot blots detect GM1 and GD1a gangliosides from homogenates of cerebral cortex and cerebellum of WT and GD3S^−/−^ mice (n = 5 per group).

### 3.5. Na+/K+-ATPase and plasma membrane Ca^2+^-ATPase activity is reduced in the cortex of GD3-deficient mice

To determine whether variations in ganglioside composition affect ATPase function, we assessed the enzyme activities of NKA and PMCA in cerebrocortical and cerebellar homogenates from GD3S^−/−^ and WT mice. In the cortex of GD3S^−/−^ mice, NKA activity was significantly lower than in WT controls (p < 0.05, unpaired two-tailed t-test) (Figure 4A). Conversely, there was no difference in the cerebellum. The same trend was observed for PMCA activity, found to be considerably lower in GD3S^−/−^ than in the WT cortex (p < 0.001, unpaired two-tailed t-test) (Figure 4D), while no change was detected for cerebellar PMCA activity.

**Figure 4.**
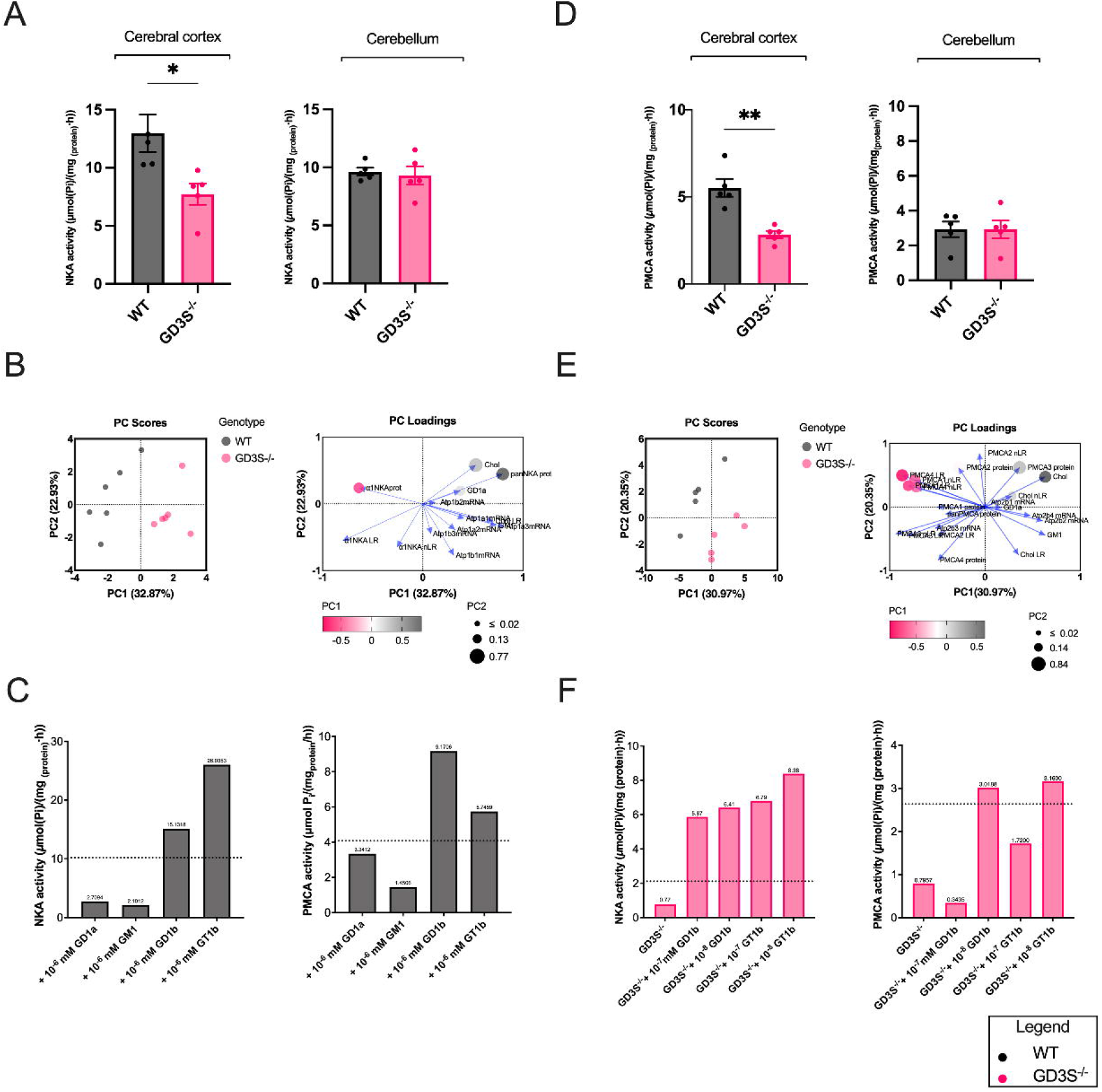
Analysis of NKA and PMCA enzyme activities in the cortex and cerebellum of GD3S^−/−^ mice. (A) NKA enzyme activity in the cerebral cortex and cerebellum of WT and GD3S^−/−^ mice. Activity was significantly reduced in the cortex of GD3S^−/−^ mice (*p < 0.05, two-tailed unpaired t-test), with no significant difference in the cerebellum. Data are presented as mean ± SEM; n = 5 mice per group. (B) Principal component analysis (PCA) of z-scored molecular features related to ATPase regulation in WT and GD3S^−/−^ mice: PCA score and loading plots for NKA-related variables. PC1 and PC2 explain 32.87% and 22.93% of the variance, respectively. (E) PCA scores and loading plots for PMCA-related variables. PC1 and PC2 explain 30.97% and 20.35% of the variance, respectively. Variables include enzyme activity, transcript and protein levels, ganglioside content, cholesterol, and the subcellular distribution of ATPases. Color reflects PC1 loading direction; circle size corresponds to PC2 contribution; WT (gray), GD3S^−/−^ (magenta), n = 3 per genotype. (D) PMCA enzyme activity in the cerebral cortex and cerebellum of WT and GD3S^−/−^ mice. Activity was significantly reduced in the cortex of GD3S^−/−^ mice (**p < 0.01, two-tailed unpaired *t*-test), with no significant difference in the cerebellum. Data are presented as mean ± SEM; n = 5 mice per group. (C) NKA and PMCA activity in WT cortical homogenates after 120-minute incubation with 10^−6^ mM of gangliosides GD1a, GM1, GD1b, or GT1b. Bars represent mean activity (stated above each bar); dotted lines indicate baseline activity without treatment. Assays were performed using pooled homogenates from 3 animals. (F) NKA and PMCA activity in GD3S^−/−^ cortical homogenates following incubation with GD1b or GT1b at 10^−7^ and 10^−8^ mM. Dotted lines indicate WT activity baseline. Mean values shown above each bar are from pooled cortical homogenates from 3 animals per group.

To determine whether GD3S deficiency leads to coordinated changes in membrane organization, we performed principal component analysis (PCA) followed by principal component regression (PCR) with enzyme activity as the outcome variable. We used centered z-score-normalized data that included ganglioside levels (GM1, GD1a), ATPase activities, cholesterol levels, and the distribution of ATPase subunits in lipid rafts (LR) and non-raft (nLR) membrane subdomains. As shown in Figures 4B and 4E, PCA clearly differentiated WT from GD3S^−/−^ mice. PCA of z-scored molecular features related to ATPase regulation in WT and GD3S^−/−^ mice (Figure 4B), namely PC scores PC1 and PC2, explains 32.87% and 22.93% of the variance, respectively. In the case of PMCA (Figure 4E), PC1 and PC2 explain 30.97% and 20.35% of the variance, respectively.

Additionally, correlation plots (Supplementary Figure 5) demonstrated associations between enzyme activities, protein expression levels, and GD1a ganglioside content in wild-type (WT) and GD3S^−/−^ mouse brains. In the cortex, NKA activity exhibited a strong positive correlation with α1 subunit protein levels and a negative correlation with overall NKA expression, indicating isoform-specific regulation. In GD3S-deficient mice, increased GD1a levels were strongly associated with decreased NKA activity (Supplementary Figure 5A). Conversely, in the cerebellum, NKA activity was unaffected by GD3S deficiency (Figure 4A), and no significant correlations were observed between NKA activity and either the α1 subunit or total catalytic α subunits expression (Supplementary Figure 5A). Although GD1a and GM1 levels were slightly higher in the GD3S^−/−^ cerebellum than in the cortex compared to WT (Figure 3E), their link to NKA activity in that brain region was minimal (Supplementary Figure 5A). PMCA activity was also strongly negatively correlated with GD1a levels (ρ < −0.7; Supplementary Figure 5B) in the cortex. Overall, these findings demonstrate that GD3S deficiency disrupts the regulation of both NKA and PMCA, primarily through alterations in membrane organization resulting from changes in ganglioside composition and the redistribution of the ion transporters.

### 3.6. Administration of exogenous b-series gangliosides restores NKA and PMCA activity

Based on previous correlation analyses and PCA visualizations, which demonstrated strong associations between GD1a and GM1 levels and changes in both NKA and PMCA activities, we conducted a screening study using cortical tissue homogenates from WT mice. Enzyme activities were assessed following administration of exogenous a-series (GM1, GD1a) and b-series (GD1b, GT1b) gangliosides at concentrations of 10^−2^, 10^−3^, 10^−4^, and 10^−6^ mM, with preincubation periods of 15, 30, 60, or 120 minutes. Control homogenates were incubated for the same duration without ganglioside treatment. Comprehensive screening data are presented in Supplementary Figure 6. Notably, the most pronounced effects were observed at lower concentrations following 120-minute preincubation. Figure 4C illustrates that GM1 and GD1a consistently inhibited NKA and PMCA activity.

Gangliosides GD1b and GT1b facilitated recovery of enzyme activity, with the extent of recovery depending on concentration (Figure 4C). The most notable recovery of ATPase activity was observed after 120-minute incubations with 10^−6^ mM GD1b or GT1b, highlighting the capacity of b-series gangliosides to modulate and restore enzyme function in GD3S deficiency, where they are not expressed. Since we observed a significant reduction in NKA activity within the cerebral cortex of GD3S^−/−^ mice and established that low concentrations of exogenous b-series gangliosides can notably increase enzymatic activity in WT mice, we further tested whether similar effects of administered b-series gangliosides could be detected in GD3S^−/−^ mice. We preincubated pooled cortical homogenates from both GD3S^−/−^ and WT mice (n = 3 per group) with GD1b and GT1b gangliosides at concentrations of 10^−7^ and 10^−8^ mM for 2 hours. This treatment resulted in a marked increase in NKA activity, approaching the level observed in untreated WT samples (Figure 4F). PMCA activity was also restored and significantly increased in GD3S^−/−^ cortical homogenates following preincubation with 10^−8^ mM GD1b and 10^−7^ and 10^−8^ mM concentrations of GT1b, compared to untreated GD3S^−/−^ controls (Figure 4F). These results indicate that reintroduction of b-series gangliosides can effectively rescue ATPase activity in neural membranes lacking endogenous b-series species due to GD3S deficiency.

## 4. Discussion

In the present study, we examined the disrupted ganglioside homeostasis following GD3S gene loss, leading to altered ion homeostasis due to changes in NKA and PMCA activity. Importantly, we showed that this impaired activity can be rescued by exogenous treatment with the gangliosides that are deficient in this mouse model.

Our initial microarray analysis (Figure 1A-B, Supplementary Table 2) was focused on the cerebellum, a brain region abundant in b-series gangliosides, where GD3S shows the highest expression in WT mice (Supplementary Figure 7). Microarray analysis provided valuable clues, specifically indicating altered expression of *Atp1b3*, which encodes a NKA subunit, and *Atp2b2*, which encodes the PMCA2 isoform (Figure 1A).

In addition, a global gene expression profile in GD3S^−/−^ brains was changed (Figure 1), which prompted the question of how membrane gangliosides could affect gene expression? Gangliosides GM1 and GD1a are overexpressed in the GD3S^−/−^ mouse model (Figure 3E, Supplementary Figure 8), which may alter gene expression. Since gangliosides selectively associate with and modulate the activity of cell-surface receptors, changes in gene expression may be downstream of cell-surface signaling (Schnaar, 2016). Alternatively, gangliosides may directly interact with transcriptomic machinery. For example, GM1 binds acetylated histones H3 and H4 at their promoters to regulate the expression of specific genes in differentiated neurons (Tsai et al., 2016). GD3 interacts with histone H1 in the nucleus (Tempera et al., 2008), and since GD3S^−/−^ mice cannot synthesize this ganglioside, gene expression may change due to increased expression of a-series and deficiency of b- and c-series gangliosides. The nuclear envelope contains other gangliosides, such as GT3, which is also absent from the brains of GD3S^−/−^ mice (Saito and Sugiyama, 2002). Therefore, various epigenetic mechanisms should be considered as contributors to altered gene expression in this and similar models, which is a promising direction for future research. Furthermore, cholesterol is known to affect the *Atp1a2* and *Atp1a3* expression via Src kinases in cell cultures (Zhang et al., 2020), and our study found that GD3S^−/−^ mice have higher cholesterol levels in lipid rafts of the cerebral cortex and cerebellum than WT mice (Figure 3C-D). Hence, altered ganglioside composition, leading to changed cholesterol levels can contribute to gene expression disturbances, pointing to a more complex mechanism of the combined effect of gangliosides and cholesterol on specific gene expression (Figure 5).

**Figure 5.**
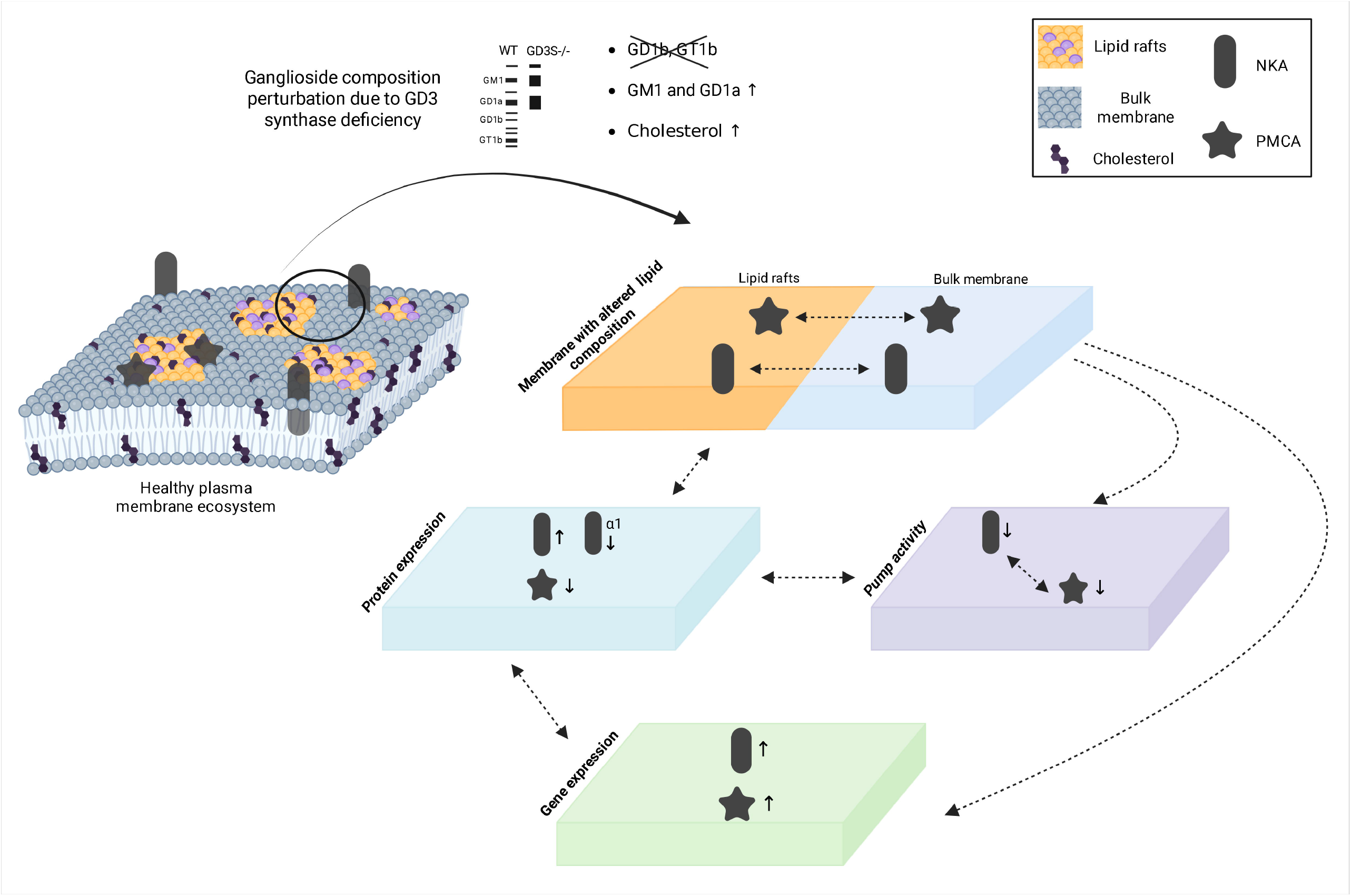
A schematic representation visualizing the potential interplay and directionality between perturbed ganglioside composition and decreased ion transporters activity. Gangliosides are primarily concentrated within lipid raft membrane microdomains. In GD3S^−/−^ mice with a markedly different ganglioside composition compared to control (WT) mice there is a redistribution of cholesterol and ion transporters NKA and PMCA, marked with bidirectional dashed arrows. This highlights the shift of these pumps between raft and bulk membrane domains (top two-color layer), influencing their activity (marked with ↓). Lower (↓) and higher (↑) gene and protein expression is also indicated for selected significant results. The altered membrane ganglioside composition and cholesterol distribution, with altered NKA/PMCA distribution (marked in top layer), can affect both NKA/PMCA activity (middle purple layer), gene expression (lower green layer), and protein expression (middle blue layer) marked with unidirectional or bidirectional dashed arrows, illustrating a top-down effect. On the other hand, differential gene expression can affect the protein expression, and *vice versa* (marked with bidirectional dashed arrow), due to transcriptional and translational regulation of NKA abundance. Bidirectional dashed arrows also connect pump activity with pump protein expression, and altered protein expression can in turn influence the positioning of the particular subunit or isoform in the plasma membrane. The featured complexity emphasizes the elaborate impact gangliosides exert on cellular homeostasis.

The distinct gene-expression pattern observed in GD3S^−/−^ brains does not translate fully to the established different protein expressions (Figure 2). Western blotting showed higher protein levels of total NKA catalytic subunits (panNKA), especially in the cortex (Figures 2A and 2C). However, when analysing α1 NKA subunit expression specifically, we observed a pattern of lower expression, in contrast to the gene expression results. There are several possible explanations for this complex relationship among varying amounts of gene and protein products for specific enzyme isoforms. These findings may be explained by the final functional consequence of these disturbances: altered enzyme activity (Figure 4A). Increased expression of the genes encoding the catalytic subunits of NKA could be mediated by a feedback loop that compensates for reduced enzyme activity by increasing enzyme production. NKA subunits are controlled extensively during development and also to accommodate physiological needs, where a combination of various transcription factors, hormones, and lipids (e.g., cholesterol and phospholipids) in conjunction with epigenetic mechanisms contribute to the regulation of NKA expression (Arystarkhova and Sweadner, 2024; Blaustein and Hamlyn, 2024; Clifford and Kaplan, 2009; Cornelius et al., 2015; Habeck et al., 2015; Li and Langhans, 2015). Our work highlights gangliosides as important additional determinants of NKA activity. Furthermore, the effects of lipids on NKA are not a one-way street: NKA can also play a role in the control of the plasma membrane cholesterol distribution (Chen et al., 2009). Because our work shows the redistribution of NKA between lipid rafts and the bulk membrane (Figure 3A), it may have an additional impact on cholesterol distribution, which in turn influences pump activity.

The most prominent finding in this work is that two ATPases, NKA and PMCA, have decreased activity in the GD3S-deficient mouse model (Figures 4A and 4D). The two pumps share an intricate connection where disturbances in ion concentration caused by one of them lead to the modified activity of the other one (Castro et al., 2006; Dimitrov, 2023; Kinoshita et al., 2022). Interestingly, in cell cultures, reaching a critically low ATP concentration induces a thermodynamic shift in free energy that favors NKA, thereby inhibiting PMCA activity. Furthermore, different gangliosides affect the two pumps differently, in a concentration-dependent manner (Figures 4C and 4F, Supplementary Figure 6), leading to decoupled regulation of their function. In addition, experiments using clickable photoaffinity ganglioside probes identified ganglioside-protein interactions between specific gangliosides and both ATPases investigated in this study (Zhang et al., 2024). Other studies delineating specific interactions between gangliosides and proteins all show that gangliosides have more interacting partners than previously thought, implying that specific binding between gangliosides and NKA/PMCA in GD3S^−/−^ mice is disrupted (Azzaz et al., 2022; Chahinian et al., 2024; Fantini et al., 2024; Fantini and Fantini, 2023). Previous reports also outlined the effect of GD3S deficiency on cholesterogenic genes expression in the brain (Mlinac et al., 2012); hence, it appears that the altered ganglioside composition in the GD3S^−/−^ model initiates a cascade of both direct effects on NKA and PMCA activity, as well as downstream indirect effects through altering cholesterol levels, lipid raft integrity, and protein and gene expression (Figure 5).

In the case of PMCA, differences in the content of specific PMCA isoforms across brain regions may indicate distinct requirements for individual isoforms with specific kinetic parameters. The results indicate that there is no simultaneous decrease in the expression of all PMCA isoforms that are expressed in the brain depending on the region, and the results are also in line with the fact that the absence of the function of neuronal-specific PMCA2 and PMCA3 in the brain cannot be compensated by increasing the content of other isoforms (Ferrington et al., 1997). A significantly reduced content of the ubiquitously expressed PMCA1 and PMCA4 isoforms, which are considered “slow” isoforms in the context of their kinetic parameters, correlates with a significant decrease in PMCA activity in the cerebral cortex tissue of the GD3S^−/−^ regions (Figure 2D and Figure 4F). However, we must keep in mind that the total number of PMCA variants resulting from alternative splicing of four genes exceeds 30 (Strehler and Zacharias, 2001). The discrepancy observed at first glance in the difference of the membrane content of individual isoforms and expression of the genes that code for them can be attributed to the fact that PMCA is considered a “long-lived” protein with a half-life of 12 (+/−1) days, but still, the half-life of individual isoforms is not known (Ferrington et al., 1997).

When examining the lipid raft integrity in GD3S^−/−^ brains, we detected a fine shift in intramembrane position in the two brain regions investigated for both ion transporters (Figure 3A-B). Not only that, but the immunoreactivity for some of the proteins (e.g., PMCA3) was found to be lower in both LRs and non-LRs in GD3S^−/−^ mice (Figure 3A-B). This effect has already been established for ganglioside-deficient mice, where the extent of this dispersion of proteins within the membrane to other membrane regions is dependent on the ganglioside environment: the more the ganglioside composition is changed, the bigger the dispersion (Ohmi et al., 2012). The interaction between PMCA, its auxiliary subunit glycoprotein neuroplastin, and specific lipids is crucial for regulating calcium transport activity and structural stability of the PMCA complex (Vinayagam et al., 2025). Our previous research showed that PMCA2 is stabilized in lipid rafts enriched for the ganglioside GM1 and neuroplastin (Ilic et al., 2021). Recent reports have demonstrated that approximately 95% of PMCA is associated with neuroplastin, a member of the immunoglobulin superfamily of cell adhesion molecules. Neuroplastin has been implicated in synaptic plasticity and is essential for cognition, long-term potentiation, and associative memory formation (Bhattacharya et al., 2017; Herrera-Molina, 2017; Lin, 2021; Lin et al., 2021; Malci et al., 2023). Hence, this finding emphasizes the importance of optimal GM1 content in the cell membrane for the stability of various protein-protein interactions.

To systematically synthesize the results obtained by examining different biological levels of expression, we performed exploratory PCA analyses (Figures 4B and 4E; Supplementary Figure 5). PCA scores clearly differentiate WT from GD3S^−/−^ mice, indicating that GD3S deficiency alters the entire molecular organization of neuronal membranes. Figure 5 schematically illustrates the complexity of ganglioside effects and highlights potential bidirectional relationships among altered ganglioside composition, cholesterol levels, gene and protein expression, and the activity of the proteins of interest. These changes and effects are mostly accentuated in the cortex; the opposite effects are also possible when comparing the cortex and cerebellum (e.g., the gene expression of PMCA isoforms, where the gene expression is higher in the cortex but lower in the cerebellum of GD3S-deficient animals, Figure 1E). This aligns with the known different spatial ganglioside architecture (Clarke et al., 2025; Lee et al., 2024), and since the investigated ion transporters are known to be regulated by various endogenous factors, these findings disclose specific gangliosides as contributors to fine-tuning NKA/PMCA activity, with differential effects in different brain regions depending on their glycolipidomic make-up.

The finding in this study with the highest translational potential is the restoration of ATPases activity by exogenously added gangliosides (Figure 4F). We performed the experiments on cortical homogenates because the cortex showed the most pronounced differences in gene and protein expression, NKA and PMCA enzyme activity, and cholesterol concentration in this work. By screening across different concentrations and incubation periods (Supplementary Figure 6), we restored enzyme activity to WT levels. The administration of exogenous gangliosides, or ganglioside replacement therapy, has already been studied for the treatment of several pathophysiological conditions (Fazzari et al., 2022; Maiti et al., 2025; Mancini et al., 2020). GD3 and GM1 intranasally regenerate neural stem cells and neurogenesis in mice with Parkinson’s disease (Fuchigami et al., 2023), and GM1 aids mouse spinal cord injury healing (Torelli et al., 2022). Ganglioside GM1, administered intravenously or subcutaneously for two years, improved motor skills and slowed disease development in two clinical trials (Schneider et al., 2013). More recently, synthetic GM1 has been shown to improve motor and memory deficits in mice with monoallelic or biallelic disruption of GM3 synthase (Chowdhury et al., 2023), underscoring the promise of this field.

## 5. Conclusion

The results of this study reveal that alterations in ganglioside composition affect ubiquitous cell membrane ion transporters, NKA and PMCA. Even though individual gene and protein expression of most subunits and isoforms do not change dramatically in GD3S^−/−^ mice, their specific expression patterns, in conjunction with altered direct ganglioside-protein interactions, lead to significantly reduced NKA and PMCA activity. Hence, the multilayer analysis demonstrates that gangliosides not only structurally organize membrane domains but also modulate the convergence of multiple regulatory tiers governing the hierarchical flow of signals that underlie optimal enzyme activity.

The lower ATPase activity observed in the GD3S-deficient mouse model is of clinical importance for two reasons: first, as more reports identify congenital disorders of ganglioside glycosylation, these findings may offer a therapeutic target for patients with deficiencies in ganglioside biosynthetic enzymes. Second, many tumor therapies target GD3S. Thus, our research on GD3S inactivation and successful restoration of ATPases activity by administering exogenous b-series ganglioside could aid in developing more specific and efficient therapies for targeting this or other enzymes in the ganglioside biosynthetic pathway.

## Abbreviations

CNS: central nervous system
CTB: cholera toxin subunit B
DEG: differentially expressed gene
GD3S: GD3 synthase
NKA: Na+/K+-ATPase
LR: lipid rafts
nLR: non-lipid rafts
PMCA: plasma membrane Ca ^2+^-ATPase receptor
WT: wild type

## Acknowledgements

We sincerely thank Prof. Ronald L. Schnaar of Johns Hopkins University for providing anti-ganglioside antibodies and for his continuous support throughout all phases of the research. We also express our gratitude to Darija Balonek-Nikolić for her help with animal handling and dissections. Figure 5 and Supplementary Figure 1 were created with BioRender.com.

## CRediT authorship contribution statement

**Borna Puljko**: Conceptualization, Validation, Formal analysis, Investigation, Visualization, Writing – original draft, Writing – review and editing. **Nikolina Maček Hrvat**: Formal analysis, Investigation, Writing – review and editing. **Katarina Ilic**: Conceptualization, Formal analysis, Investigation, Writing – review and editing. **Ana Ujevic**: Formal analysis, Investigation, Writing – review and editing. **Eva Josic**: Formal analysis, Investigation, Writing – review and editing. **Mario Stojanovic**: Formal analysis, Investigation, Writing – review and editing. **Tadeja Režen**: Formal analysis, Investigation, Visualization, Writing – review and editing. **Klementina Fon Tacer**: Formal analysis, Investigation, Visualization, Writing – review and editing. **Damjana Rozman**: Formal analysis, Funding acquisition, Resources, Supervision, Writing - Review & Editing. **Marta Balog**: Investigation, Methodology, Writing – review and editing. **Marija Heffer**: Methodology, Funding acquisition, Resources, Supervision, Writing - Review & Editing. **Svjetlana Kalanj-Bognar**: Conceptualization, Funding acquisition, Project administration, Resources, Supervision, Writing - Review & Editing. **Kristina Mlinac-Jerkovic**: Conceptualization, Validation, Formal analysis, Funding acquisition, Investigation, Project administration, Resources, Supervision, Visualization, Writing – original draft, Writing – review and editing

## Consent for publication

All authors read and approved the final manuscript.

## Ethics approval

All animal procedures were approved by the relevant institutional and national authorities (Ethics Committee of the University of Zagreb for the Croatian Science Foundation projects IP-2016-06-8636 to S.K.-B., IP-2014-09-2324 to M.H.) and complied with relevant animal welfare regulations (Animal Protection Act 102/2017; Ordinance 55/2013) and EU Directive 2010/63/EU., including ARRIVE 2.0 guidelines and the 3R principles.

## References

Abreu, C.A., Teixeira-Pinheiro, L.C., Lani-Louzada, R., da Silva-Junior, A.J., Vasques, J.F., Gubert, F., Nascimento-dos-Santos, G., Mohana-Borges, R., Matos, E. de S., Pimentel-Coelho, P.M., Santiago, M.F., Mendez-Otero, R., 2021. GD3 synthase deletion alters retinal structure and impairs visual function in mice. J. Neurochem. 158, 694–709.

Anand, V., El-Dana, F., Baran, N., Borgman, J., Yin, Z., Zhao, H., Wong, S.T., Andreeff, M., Battula, V.L., 2025. GD3 synthase drives resistance to p53-induced apoptosis in breast cancer by modulating mitochondrial function. Oncogene 44, 2646–2661.

Arystarkhova, E., Sweadner, K.J., 2024. Na,K-ATPase expression can be limited post-transcriptionally: A test of the role of the beta subunit, and a review of evidence. Int. J. Mol. Sci. 25, 7414.

Azzaz, F., Yahi, N., Di Scala, C., Chahinian, H., Fantini, J., 2022. Ganglioside binding domains in proteins: Physiological and pathological mechanisms. Adv. Protein Chem. Struct. Biol. 128, 289–324.

Benarroch, E.E., 2011. Na+, K+-ATPase: functions in the nervous system and involvement in neurologic disease. Neurology 76, 287–293.

Bernardo, A., Harrison, F.E., McCord, M., Zhao, J., Bruchey, A., Davies, S.S., Jackson Roberts, L., Mathews, P.M., Matsuoka, Y., Ariga, T., Yu, R.K., Thompson, R., McDonald, M.P., 2009. Elimination of GD3 synthase improves memory and reduces amyloid-β plaque load in transgenic mice. Neurobiol. Aging 30, 1777–1791.

Berrocal, M., Mata, A.M., 2023. The plasma membrane Ca2+-ATPase, a molecular target for tau-induced cytosolic calcium dysregulation. Neuroscience 518, 112–118.

Bhattacharya, S., Herrera-Molina, R., Sabanov, V., Ahmed, T., Iscru, E., Stöber, F., Richter, K., Fischer, K.-D., Angenstein, F., Goldschmidt, J., Beesley, P.W., Balschun, D., Smalla, K.-H., Gundelfinger, E.D., Montag, D., 2017. Genetically induced retrograde amnesia of associative memories after neuroplastin ablation. Biol. Psychiatry 81, 124–135.

Blaustein, M.P., Hamlyn, J.M., 2024. Sensational site: the sodium pump ouabain-binding site and its ligands. Am. J. Physiol. Cell Physiol. 326, C–C1177.

Boczek, T., Radzik, T., Ferenc, B., Zylinska, L., 2019. The Puzzling Role of Neuron-Specific PMCA Isoforms in the Aging Process. Int. J. Mol. Sci. 20. 10.3390/IJMS20246338

Bookout, A.L., Cummins, C.L., Mangelsdorf, D.J., Pesola, J.M., Kramer, M.F., 2006. High-throughput real-time quantitative reverse transcription PCR. Curr. Protoc. Mol. Biol. Chapter 15, Unit 15.8.

Bøttger, P., Doĝanli, C., Lykke-Hartmann, K., 2012. Migraine- and dystonia-related disease-mutations of Na+/K+-ATPases: Relevance of behavioral studies in mice to disease symptoms and neurological manifestations in humans. Neurosci. Biobehav. Rev. 36, 855–871.

Bøttger, P., Tracz, Z., Heuck, A., Nissen, P., Romero-Ramos, M., Lykke-Hartmann, K., 2011. Distribution of Na/K-ATPase alpha 3 isoform, a sodium-potassium P-type pump associated with rapid-onset of dystonia parkinsonism (RDP) in the adult mouse brain. J. Comp. Neurol. 519, 376–404.

Cao, S., Hu, X., Ren, S., Wang, Y., Shao, Y., Wu, K., Yang, Z., Yang, W., He, G., Li, X., 2023. The biological role and immunotherapy of gangliosides and GD3 synthase in cancers. Frontiers in Cell and Developmental Biology 11, 113.

Castro, J., Ruminot, I., Porras, O.H., Flores, C.M., Hermosilla, T., Verdugo, E., Venegas, F., Härtel, S., Michea, L., Barros, L.F., 2006. ATP steal between cation pumps: a mechanism linking Na+ influx to the onset of necrotic Ca2+ overload. Cell Death Differ. 13, 1675–1685.

Chahinian, H., Yahi, N., Fantini, J., 2024. Glutamate, gangliosides, and the synapse: Electrostatics at work in the brain. Int. J. Mol. Sci. 25, 8583.

Chen, E.Y., Tan, C.M., Kou, Y., Duan, Q., Wang, Z., Meirelles, G.V., Clark, N.R., Ma’ayan, A., 2013. Enrichr: interactive and collaborative HTML5 gene list enrichment analysis tool. BMC Bioinformatics 14. 10.1186/1471-2105-14-128

Chen, Y., Cai, T., Wang, H., Li, Z., Loreaux, E., Lingrel, J.B., Xie, Z., 2009. Regulation of intracellular cholesterol distribution by Na/K-ATPase. J. Biol. Chem. 284, 14881–14890.

Chowdhury, S., Kumar, R., Zepeda, E., DeFrees, S., Ledeen, R., 2023. Synthetic GM1 improves motor and memory dysfunctions in mice with monoallelic or biallelic disruption of GM3 synthase. FEBS Open Bio. 10.1002/2211-5463.13669

Clarke, H.A., Ma, X., Shedlock, C.J., Medina, T., Hawkinson, T.R., Wu, L., Ribas, R.A., Keohane, S., Ravi, S., Bizon, J.L., Burke, S.N., Abisambra, J.F., Merritt, M.E., Prentice, B.M., Vander Kooi, C.W., Gentry, M.S., Chen, L., Sun, R.C., 2025. Spatial mapping of the brain metabolome lipidome and glycome. Nat. Commun. 16, 4373.

Clifford, R.J., Kaplan, J.H., 2009. Regulation of Na,K-ATPase subunit abundance by translational repression. J. Biol. Chem. 284, 22905–22915.

Cohen, M., Varki, A., 2010. The sialome-far more than the sum of its parts. OMICS 14, 455–464.

Cornelius, F., Habeck, M., Kanai, R., Toyoshima, C., Karlish, S.J.D., 2015. General and specific lipid–protein interactions in Na,K-ATPase. Biochimica et Biophysica Acta (BBA) - Biomembranes 1848, 1729–1743.

Dimitrov, A.G., 2023. Resting membrane state as an interplay of electrogenic transporters with various pumps. Pflugers Arch. 475, 1113–1128.

Domi, T., Di Leva, F., Fedrizzi, L., Rimessi, A., Brini, M., 2007. Functional specificity of PMCA isoforms? Ann. N. Y. Acad. Sci. 1099, 237–246.

Fantini, J., Azzaz, F., Bennaï, R., Yahi, N., Chahinian, H., 2024. Cholesterol-dependent serotonin insertion controlled by gangliosides in model lipid membranes. Int. J. Mol. Sci. 25. 10.3390/ijms251810194

Fantini, J., Fantini, C.J., 2023. Lipid rafts and human diseases: why we need to target gangliosides. FEBS Open Bio 13, 1636–1650.

Fazzari, M., Lunghi, G., Chiricozzi, E., Mauri, L., Sonnino, S., 2022. Gangliosides and the Treatment of Neurodegenerative Diseases: A Long Italian Tradition. Biomedicines 10. 10.3390/BIOMEDICINES10020363

Ferrington, D.A., Chen, X., Krainev, A.G., Michaelis, E.K., Bigelow, D.J., 1997. Protein half-lives of calmodulin and the plasma membrane Ca-ATPase in rat brain. Biochem. Biophys. Res. Commun. 237, 163–165.

Fuchigami, T., Itokazu, Y., Morgan, J.C., Yu, R.K., 2023. Restoration of Adult Neurogenesis by Intranasal Administration of Gangliosides GD3 and GM1 in The Olfactory Bulb of A53T Alpha-Synuclein-Expressing Parkinson’s-Disease Model Mice. Mol. Neurobiol. 10.1007/S12035-023-03282-2

Ge, S.X., Jung, D., Yao, R., 2020. ShinyGO: a graphical gene-set enrichment tool for animals and plants. Bioinformatics 36, 2628–2629.

Groux-Degroote, S., Guérardel, Y., Delannoy, P., 2017. Gangliosides: Structures, Biosynthesis, Analysis, and Roles in Cancer. Chembiochem 18, 1146–1154.

Gu, H.-O., Noh, S.W., Kim, O.-H., Oh, B.-C., 2025. Crucial roles of calcium ATPases and phosphoinositides: Insights into pathophysiology and therapeutic strategies. Mol. Cells 48, 100254.

Habeck, M., Haviv, H., Katz, A., Kapri-Pardes, E., Ayciriex, S., Shevchenko, A., Ogawa, H., Toyoshima, C., Karlish, S.J.D., 2015. Stimulation, Inhibition, or Stabilization of Na,K-ATPase Caused by Specific Lipid Interactions at Distinct Sites. J. Biol. Chem. 290, 4829.

Handa, Y., Ozaki, N., Honda, T., Furukawa, Koichi, Tomita, Y., Inoue, M., Furukawa, Keiko, Okada, M., Sugiura, Y., 2005. GD3 synthase gene knockout mice exhibit thermal hyperalgesia and mechanical allodynia but decreased response to formalin-induced prolonged noxious stimulation. Pain 117, 271–279.

Herrera-Molina, R., 2017. Deafness causing neuroplastin missense variants fail to promote plasma membrane Ca2+-ATPase levels and Ca2+ transient regulation in brain neurons. Neuroplastin deletion in glutamatergic neurons impairs selective brain functions and calcium regulation: Implication for cognitive deterioration. Sci. Rep 7, 1705–1706.

Husain, S., Yildirim-Toruner, C., Rubio, J.P., Field, J., Schwalb, M., Cook, S., Devoto, M., Vitale, E., 2008. Variants of ST8SIA1 are associated with risk of developing multiple sclerosis. PLoS One 3. 10.1371/JOURNAL.PONE.0002653

Ilic, K., Lin, X., Malci, A., Stojanović, M., Puljko, B., Rožman, M., Vukelić, Ž., Heffer, M., Montag, D., Schnaar, R.L., Kalanj-Bognar, S., Herrera-Molina, R., Mlinac-Jerkovic, K., 2021. Plasma Membrane Calcium ATPase-Neuroplastin Complexes Are Selectively Stabilized in GM1-Containing Lipid Rafts. Int. J. Mol. Sci. 22, 13590.

Jiang, L., Bechtel, M.D., Bean, J.L., Winefield, R., Williams, T.D., Zaidi, A., Michaelis, E.K., Michaelis, M.L., 2014. Effects of gangliosides on the activity of the plasma membrane Ca 2 +-ATPase. Biochimica et Biophysica Acta - Biomembranes 1838, 1255–1265.

Kaplan, J.H., 2002. Biochemistry of Na,K-ATPase. Annu. Rev. Biochem. 71, 511–535.

Kasprowicz, A., Sophie, G.D., Lagadec, C., Delannoy, P., 2022. Role of GD3 Synthase ST8Sia I in Cancers. Cancers 2022, Vol. 14, Page 1299 14, 1299.

Kinoshita, P.F., Orellana, A.M.M., Nakao, V.W., de Souza Port’s, N.M., Quintas, L.E.M., Kawamoto, E.M., Scavone, C., 2022. The Janus face of ouabain in Na+ /K+-ATPase and calcium signalling in neurons. Br. J. Pharmacol. 179, 1512–1524.

Kip, S.N., Strehler, E.E., 2003. Characterization of PMCA isoforms and their contribution to transcellular Ca2+ flux in MDCK cells. American Journal of Physiology - Renal Physiology 284, 122–132.

Kittaka, D., Itoh, M.I., Ohmi, Y., Kondo, Y., Fukumoto, S., Urano, T., Tajima, O., Furukawa, Keiko, Furukawa, Koichi, 2008. Impaired hypoglossal nerve regeneration in mutant mice lacking complex gangliosides: Down-regulation of neurotrophic factors and receptors as possible mechanisms. Glycobiology 18, 509–516.

Krebs, J., 2015. The plethora of PMCA isoforms: Alternative splicing and differential expression. Biochimica et Biophysica Acta (BBA) - Molecular Cell Research 1853, 2018–2024.

Kuleshov, M.V., Jones, M.R., Rouillard, A.D., Fernandez, N.F., Duan, Q., Wang, Z., Koplev, S., Jenkins, S.L., Jagodnik, K.M., Lachmann, A., Mcdermott, M.G., Monteiro, C.D., Gundersen, G.W., Ma’ayan, A., 2016. Enrichr: a comprehensive gene set enrichment analysis web server 2016 update. Nucleic Acids Research.

Lee, J., Yin, D., Yun, J., Kim, M., Kim, S.-W., Hwang, H., Park, J.E., Lee, B., Lee, C.J., Shin, H.-S., An, H.J., 2024. Deciphering mouse brain spatial diversity via glyco-lipidomic mapping. Nat. Commun. 15. 10.1038/s41467-024-53032-8

Leon, A., Facci, L., Toffano, G., Sonnino, S., Tettamanti, G., 1981. Activation of (Na+, K+)-ATPase by Nanomolar Concentrations of GM1 Ganglioside. J. Neurochem. 37, 350–357.

Li, Z., Langhans, S.A., 2015. Transcriptional regulators of Na,K-ATPase subunits. Front. Cell Dev. Biol. 3, 66.

Lin, X., 2021. Neuroplastin expression is essential for hearing and hair cell PMCA expression. Brain Struct. Funct 226, 1533–1551.

Lin, X., Liang, Y., Herrera-Molina, R., Montag, D., 2021. Neuroplastin in neuropsychiatric diseases. Genes (Basel) 12, 1507.

Liu, J., Zheng, X., Pang, X., Li, L., Wang, J., Yang, C., Du, G., 2018. Ganglioside GD3 synthase (GD3S), a novel cancer drug target. Yao Xue Xue Bao 8, 713.

Lorent, J.H., Diaz-Rohrer, B., Lin, X., Spring, K., Gorfe, A.A., Levental, K.R., Levental, I., 2017. Structural determinants and functional consequences of protein affinity for membrane rafts. Nat. Commun. 8, 1219.

Maiti, P., Xue, Y., Rex, T.S., McDonald, M.P., 2025. Gene therapy targeting GD3 synthase protects against MPTP-induced parkinsonism and executive dysfunction. Eur. J. Neurosci. 61, e70061.

Malci, A., Lin, X., Shi, Y.S., Herrera-Molina, R., 2023. Neuroplastin in Ca2+ signal regulation and plasticity of glutamatergic synapses. Neural Regen. Res. 18, 1705–1706.

Maloy, A., Alexander, S., Andreas, A., Nyunoya, T., Chandra, D., 2022. Stain-Free total-protein normalization enhances the reproducibility of Western blot data. Anal. Biochem. 654, 114840.

Mancini, G., Loberto, N., Olioso, D., Dechecchi, M.C., Cabrini, G., Mauri, L., Bassi, R., Schiumarini, D., Chiricozzi, E., Lippi, G., Pesce, E., Sonnino, S., Pedemonte, N., Tamanini, A., Aureli, M., 2020. GM1 as Adjuvant of Innovative Therapies for Cystic Fibrosis Disease. International Journal of Molecular Sciences 2020, Vol. 21, Page 4486 21, 4486.

Mlinac, K., Fabris, D., Vukelić, Z., Rožman, M., Heffer, M., Bognar, S.K., 2013. Structural analysis of brain ganglioside acetylation patterns in mice with altered ganglioside biosynthesis. Carbohydr. Res. 382, 1–8.

Mlinac, K., Fon Tacer, K., Heffer, M., Rozman, D., Kalanj Bognar, S., 2012. Cholesterogenic genes expression in brain and liver of ganglioside-deficient mice. Mol. Cell. Biochem. 369, 127–133.

Mlinac-Jerkovic, K., Heffer, M., Schnaar, R.L., 2026. Gangliosides in molecular interactions and cell regulation. J. Biol. Chem. 111184.

Mlinac-Jerkovic, K., Ilic, K., Zjalić, M., Mandić, D., Debeljak, Ž., Balog, M., Damjanović, V., Maček Hrvat, N., Habek, N., Kalanj-Bognar, S., Schnaar, R.L., Heffer, M., 2021. Who’s in, who’s out? Re-evaluation of lipid raft residents. J. Neurochem. jnc.15446.

Ohkawa, Y., Zhang, P., Momota, H., Kato, A., Hashimoto, N., Ohmi, Y., Bhuiyan, R.H., Farhana, Y., Natsume, A., Wakabayashi, T., Furukawa, Keiko, Furukawa, Koichi, 2021. Lack of GD3 synthase (St8sia1) attenuates malignant properties of gliomas in genetically engineered mouse model. Cancer Sci. 112, 3756.

Ohmi, Y., Ohkawa, Y., Yamauchi, Y., Tajima, O., Furukawa, Keiko, Furukawa, Koichi, 2012. Essential roles of gangliosides in the formation and maintenance of membrane microdomains in brain tissues. Neurochem. Res. 37, 1185–1191.

Orr, S.L., Le, D., Long, J.M., Sobieszczuk, P., Ma, B., Tian, H., Fang, X., Paulson, J.C., Marth, J.D., Varki, N., 2013. A phenotype survey of 36 mutant mouse strains with gene-targeted defects in glycosyltransferases or glycan-binding proteins. Glycobiology 23, 363–380.

Persaud-Sawin, D.-A., Lightcap, S., Harry, G.J., 2009. Isolation of rafts from mouse brain tissue by a detergent-free method. J. Lipid Res. 50, 759–767.

Pietrobon, D., Conti, F., 2024. Astrocytic Na+, K+ ATPases in physiology and pathophysiology. Cell Calcium 118, 102851.

Pirahanchi, Y., Jessu, R., Aeddula, N.R., 2021. Physiology, Sodium Potassium Pump. StatPearls.

Puljko, B., Grbavac, J., Potočki, V., Ilic, K., Viljetić, B., Kalanj-Bognar, S., Heffer, M., Debeljak, Ž., Blažetić, S., Mlinac-Jerkovic, K., 2024. The good, the bad, and the unknown nature of decreased GD3 synthase expression. Front. Mol. Neurosci. 17, 1465013.

Puljko, B., Stojanović, M., Ilic, K., Maček Hrvat, N., Zovko, A., Damjanović, V., Mlinac-Jerkovic, K., Kalanj-Bognar, S., 2021. Redistribution of gangliosides accompanies thermally induced Na+, K+-ATPase activity alternation and submembrane localisation in mouse brain. Biochimica et Biophysica Acta (BBA) - Biomembranes 1863, 183475.

Ramagopalan, S.V., Morrison, K.M., Para, A., Handel, A., Disanto, G., Handunnetthi, L., Orton, S.M., Sadovnick, A.D., Ebers, G.C., 2009. Variants in ST8SIA1 do not play a major role in susceptibility to multiple sclerosis in Canadian families. J. Neuroimmunol. 212, 142–144.

Ribeiro-Resende, V.T., Araújo Gomes, T., de Lima, S., Nascimento-Lima, M., Bargas-Rega, M., Santiago, M.F., Reis, R.A. de M., de Mello, F.G., 2014. Mice lacking GD3 synthase display morphological abnormalities in the sciatic nerve and neuronal disturbances during peripheral nerve regeneration. PLoS One 9, e108919.

Rivero-Gutiérrez, B., Anzola, A., Martínez-Augustin, O., de Medina, F.S., 2014. Stain-free detection as loading control alternative to Ponceau and housekeeping protein immunodetection in Western blotting. Anal. Biochem. 467, 1–3.

Saito, M., Sugiyama, K., 2002. Characterization of nuclear gangliosides in rat brain: Concentration, composition, and developmental changes. Arch. Biochem. Biophys. 398, 153–159.

Schnaar, R.L., 2019. The Biology of Gangliosides. Adv. Carbohydr. Chem. Biochem. 76, 113–148.

Schnaar, R.L., 2016. Gangliosides of the vertebrate nervous system. J. Mol. Biol. 428, 3325–3336.

Schneider, J.S., Gollomp, S.M., Sendek, S., Colcher, A., Cambi, F., Du, W., 2013. A Randomized, Controlled, Delayed Start Trial of GM1 Ganglioside in Treated Parkinson’s Disease Patients. J. Neurol. Sci. 324, 140.

Sipione, S., Monyror, J., Galleguillos, D., Steinberg, N., Kadam, V., 2020. Gangliosides in the brain: Physiology, pathophysiology and therapeutic applications. Front. Neurosci. 14. 10.3389/fnins.2020.572965

Strehler, E.E., Zacharias, D.A., 2001. Role of alternative splicing in generating isoform diversity among plasma membrane calcium pumps. Physiol. Rev. 81, 21–50.

Sun, J., Zheng, Y., Chen, Z., Wang, Y., 2022. The role of Na+-K+-ATPase in the epileptic brain. CNS Neurosci. Ther. 28, 1294–1302.

Svennerholm, L., Boström, K., Fredman, P., Månsson, J.E., Rosengren, B., Rynmark, B.M., 1989. Human brain gangliosides: developmental changes from early fetal stage to advanced age. Biochim. Biophys. Acta 1005, 109–117.

Tang, F.L., Wang, J., Itokazu, Y., Yu, R.K., 2021. Ganglioside GD3 regulates dendritic growth in newborn neurons in adult mouse hippocampus via modulation of mitochondrial dynamics. J. Neurochem. 156, 819–833.

Tempera, I., Buchetti, B., Lococo, E., Gradini, R., Mastronardi, A., Mascellino, M.T., Sale, P., Mosca, L., d’Erme, M., Lenti, L., 2008. GD3 nuclear localization after apoptosis induction in HUT-78 cells. Biochem. Biophys. Res. Commun. 368, 495–500.

Torelli, A.G., Cristante, A.F., de Barros-Filho, T.E.P., dos Santos, G.B., Morena, B.C., Correia, F.F., Paschon, V., 2022. Effects of ganglioside GM1 and erythropoietin on spinal cord injury in mice: Functional and immunohistochemical assessments. Clinics 77. 10.1016/J.CLINSP.2022.100006

Tsai, Y.-T., Itokazu, Y., Yu, R.K., 2016. GM1 ganglioside is involved in epigenetic activation loci of neuronal cells. Neurochem. Res. 41, 107–115.

Vinayagam, D., Sitsel, O., Schulte, U., Constantin, C.E., Oosterheert, W., Prumbaum, D., Zolles, G., Fakler, B., Raunser, S., 2025. Molecular mechanism of ultrafast transport by plasma membrane Ca2+-ATPases. Nature 646, 236–245.

Wang, J., Cheng, A., Wakade, C., Yu, R.K., 2014. Ganglioside GD3 is required for neurogenesis and long-term maintenance of neural stem cells in the postnatal mouse brain. J. Neurosci. 34, 13790–13800.

Watanabe, Y., Nara, K., Takahashi, H., Nagai, Y., Sanai, Y., 1996. The molecular cloning and expression of alpha 2,8-sialyltransferase (GD3 synthase) in a rat brain. J. Biochem. 120, 1020–1027.

Xie, Z., Bailey, A., Kuleshov, M.V., Clarke, D.J.B., Evangelista, J.E., Jenkins, S.L., Lachmann, A., Wojciechowicz, M.L., Kropiwnicki, E., Jagodnik, K.M., Jeon, M., Ma’ayan, A., 2021. Gene set knowledge discovery with Enrichr. Curr. Protoc. 1, e90.

Yamaguchi, T., Kato, K., 2018. Molecular Dynamics of Gangliosides, in: Methods in Molecular Biology, Methods in Molecular Biology (Clifton, N.J.). Springer New York, New York, NY, pp. 411–417.

Yamamoto, A., Haraguchi, M., Yamashiro, S., Fukumoto, S., Furukawa, K., Takamiya, K., Atsuta, M., Shiku, H., Furukawa, K., 1996. Heterogeneity in the expression pattern of two ganglioside synthase genes during mouse brain development. J. Neurochem. 66, 26–34.

Ye, J., Coulouris, G., Zaretskaya, I., Cutcutache, I., Rozen, S., Madden, T.L., 2012. Primer-BLAST: a tool to design target-specific primers for polymerase chain reaction. BMC Bioinformatics 13, 134.

Ygberg, S., Akkuratov, E.E., Howard, R.J., Taylan, F., Jans, D.C., Mahato, D.R., Katz, A., Kinoshita, P.F., Portal, B., Nennesmo, I., Lindskog, M., Karlish, S.J.D., Andersson, M., Lindstrand, A., Brismar, H., Aperia, A., 2021. A missense mutation converts the Na+,K+-ATPase into an ion channel and causes therapy-resistant epilepsy. J. Biol. Chem. 297. 10.1016/J.JBC.2021.101355/ATTACHMENT/B34524EF-49A9-45E6-9A77-9F5811DFCFC4/MMC1.PDF

Yu, R.K., Tsai, Y.T., Ariga, T., Yanagisawa, M., 2011. Structures, biosynthesis, and functions of gangliosides—An overview. J. Oleo Sci. 60, 537.

Zhang, G.-L., Porter, M.J., Awol, A.K., Orsburn, B.C., Canner, S.W., Gray, J.J., O’Meally, R.N., Cole, R.N., Schnaar, R.L., 2024. The Human Ganglioside Interactome in Live Cells Revealed Using Clickable Photoaffinity Ganglioside Probes. J. Am. Chem. Soc. 146, 17801–17816.

Zhang, J., Li, X., Yu, H., Larre, I., Dube, P.R., Kennedy, D.J., Wilson Tang, W.H., Westfall, K., Pierre, S.V., Xie, Z., Chen, Y., 2020. Regulation of Na/K-ATPase expression by cholesterol: isoform specificity and the molecular mechanism. American Journal of Physiology - Cell Physiology 319, C1107.

Zhang, P., Ohkawa, Y., Yamamoto, S., Momota, H., Kato, A., Kaneko, K., Natsume, A., Farhana, Y., Ohmi, Y., Okajima, T., Bhuiyan, R.H., Wakabayashi, T., Furukawa, Keiko, Furukawa, Koichi, 2021. St8sia1-deficiency in mice alters tumor environments of gliomas, leading to reduced disease severity. Nagoya J. Med. Sci. 83, 535–549.

