## Supplementary material for "GD3 synthase deficiency disrupts Na^+^/K^+^-ATPase and plasma membrane Ca^2+^-ATPase function in mouse brain"

### SUPPLEMENTARY FIGURES

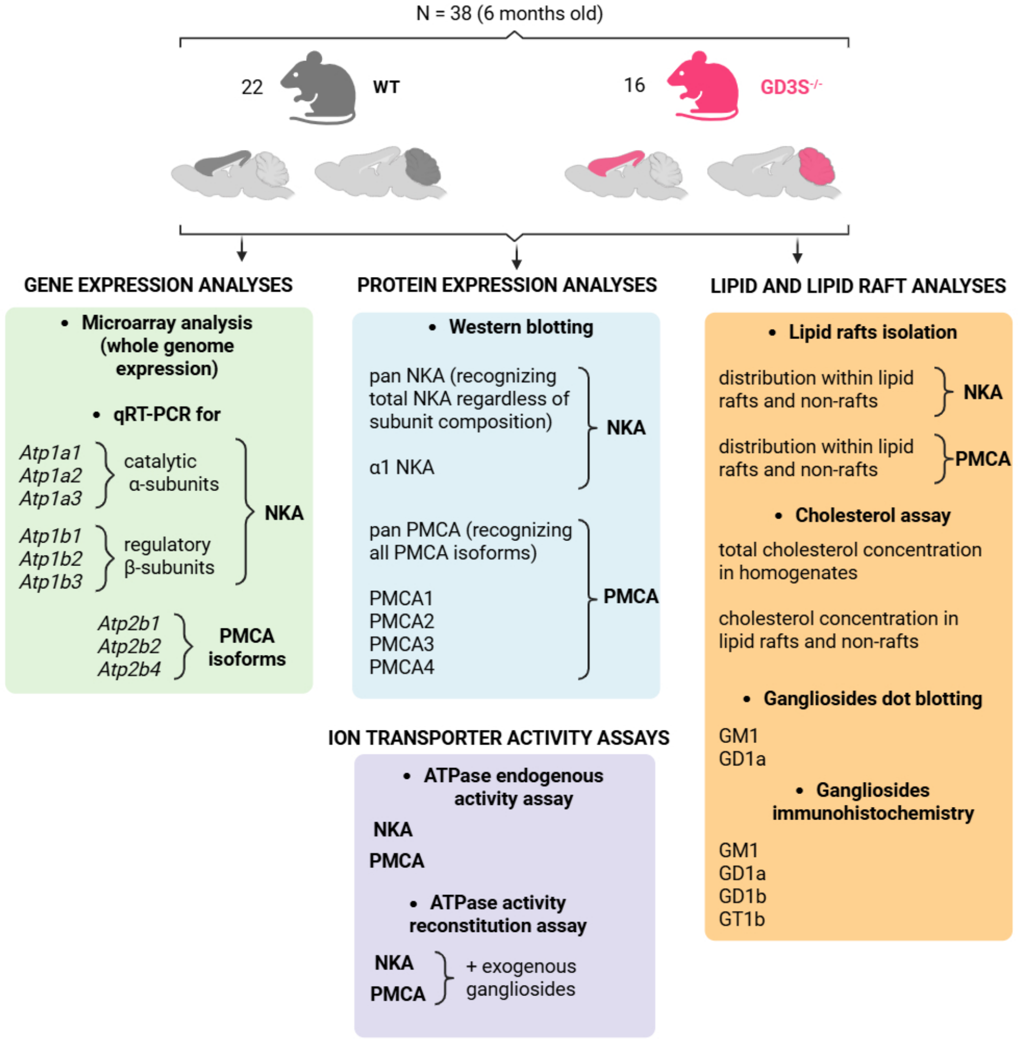


**Supplementary Figure 1.** An overview of the study and the performed analyses.

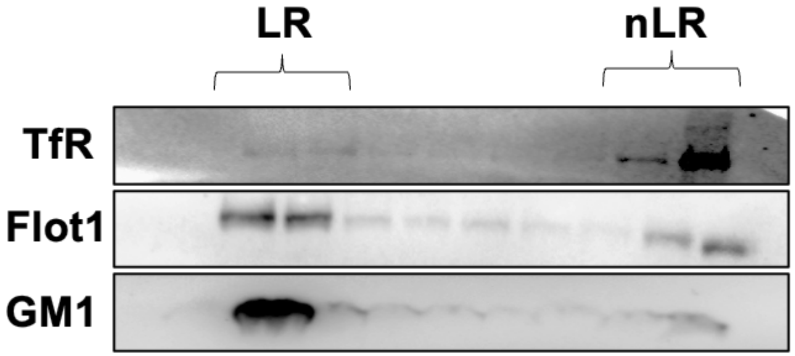


**Supplementary Figure 2.** Validation of lipid raft (LR) and non-lipid raft (nLR) isolation success by Western blotting.
Lipid raft fractions were validated by enrichment of established LR markers flotillin-1 (Flot1) and the ganglioside GM1, the latter detected by cholera toxin B subunit (CTB). Transferrin receptor (TfR), a marker of non-raft membranes, was enriched in non-lipid raft (nLR) fractions. Representative blots are shown.

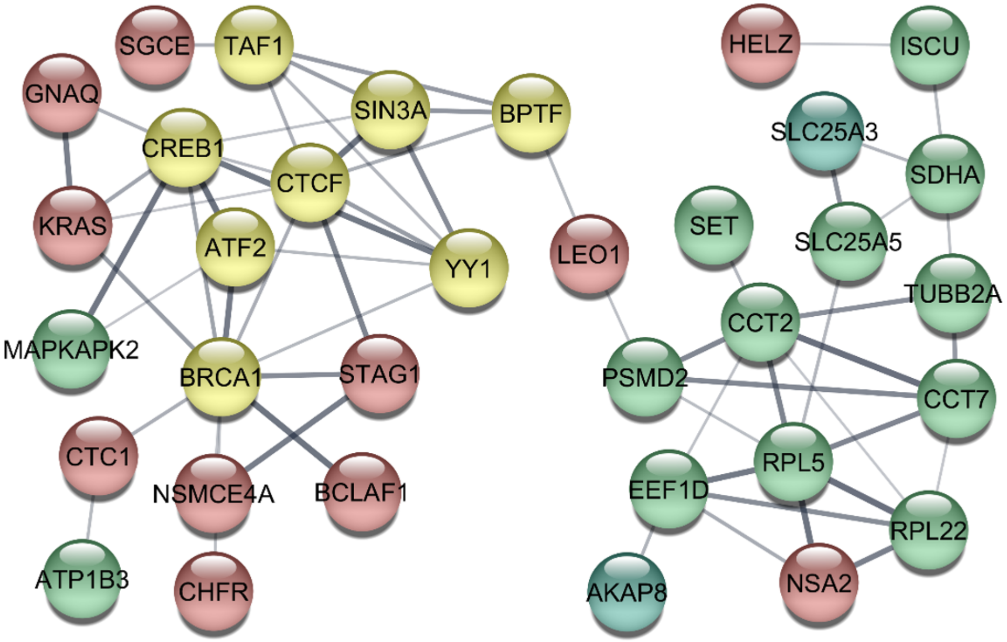


**Supplementary Figure 3.** Network-like interactions between DEG (green - downregulated, red – upregulated) and selected enriched transcription factors (yellow). Cytoscape analysis using STRING database of DEG and selected significantly enriched transcription factors by Enricher in ENCODE and ChEA databases (TAF1, ATF2, BRCA1, CREB1, CTCF, YY1, SIN3A) and TRANSFAC analysis of DEG promotors (BPTF).

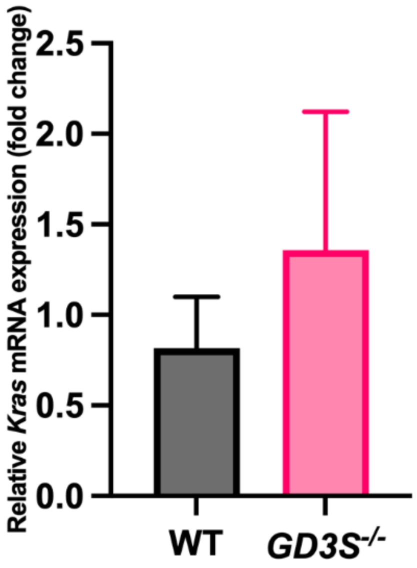


**Supplementary Figure 4.** Gene expression of *Kras* in cerebellar tissue of WT and GD3S$^{-/-}$ mice analyzed by qRT-PCR. Bars show group mean ± SEM (n = 3 animals/group), expressed as fold change from the control means.

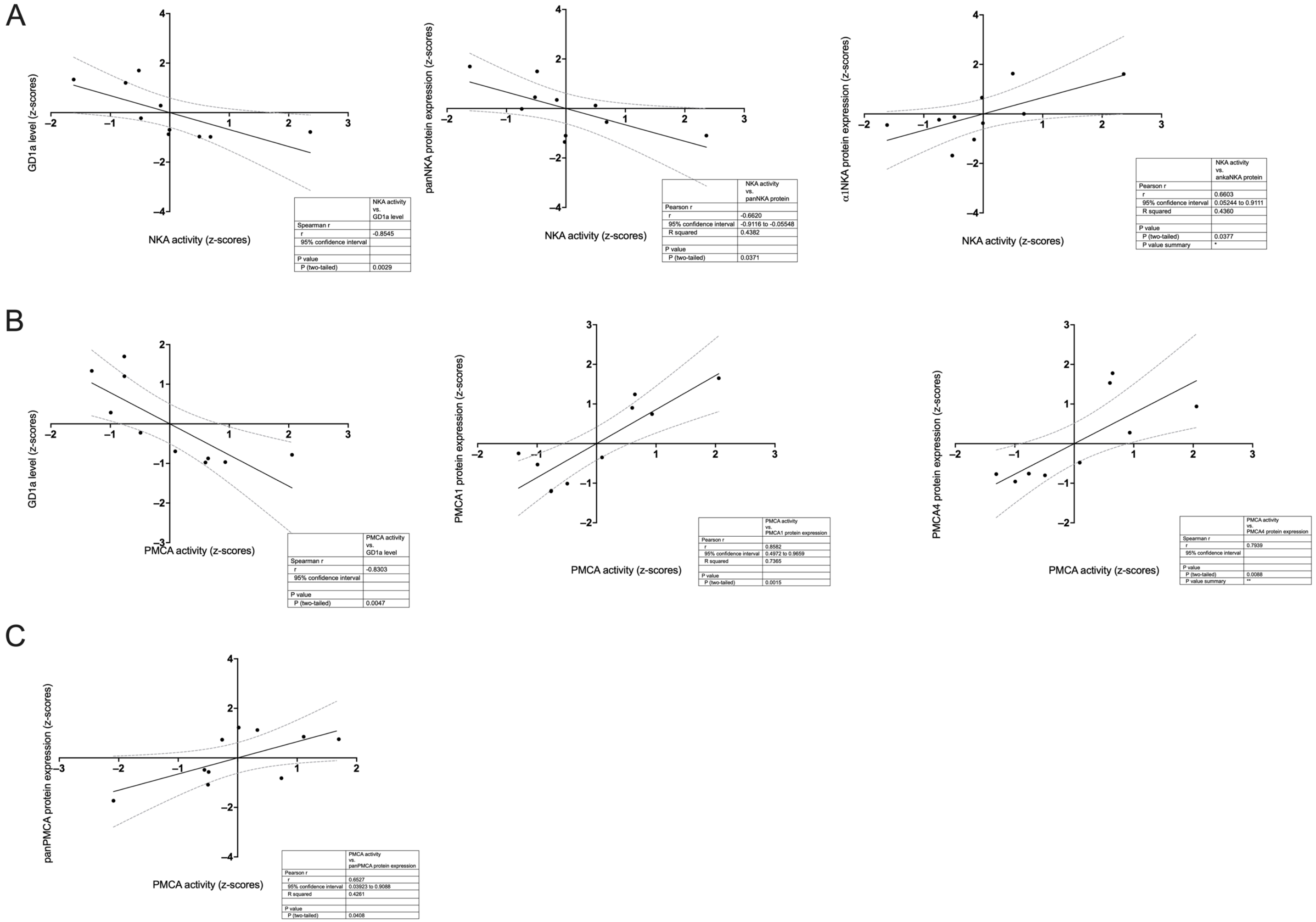


**Supplementary Figure 5.** Correlation plots demonstrate associations between enzyme activities, protein expression levels, and GD1a ganglioside content in wild-type (WT) and GD3S⁻/⁻ mouse brains. (A) In the cerebral cortex, significant correlations were observed between NKA activity and GD1a levels, pan-NKA, and α1-NKA expression. (B) In the cortex, PMCA activity correlated with GD1a levels and with the expression of PMCA1 and PMCA4. (C) In the cerebellum, PMCA activity correlated with pan-PMCA expression. All variables are z-score normalized.

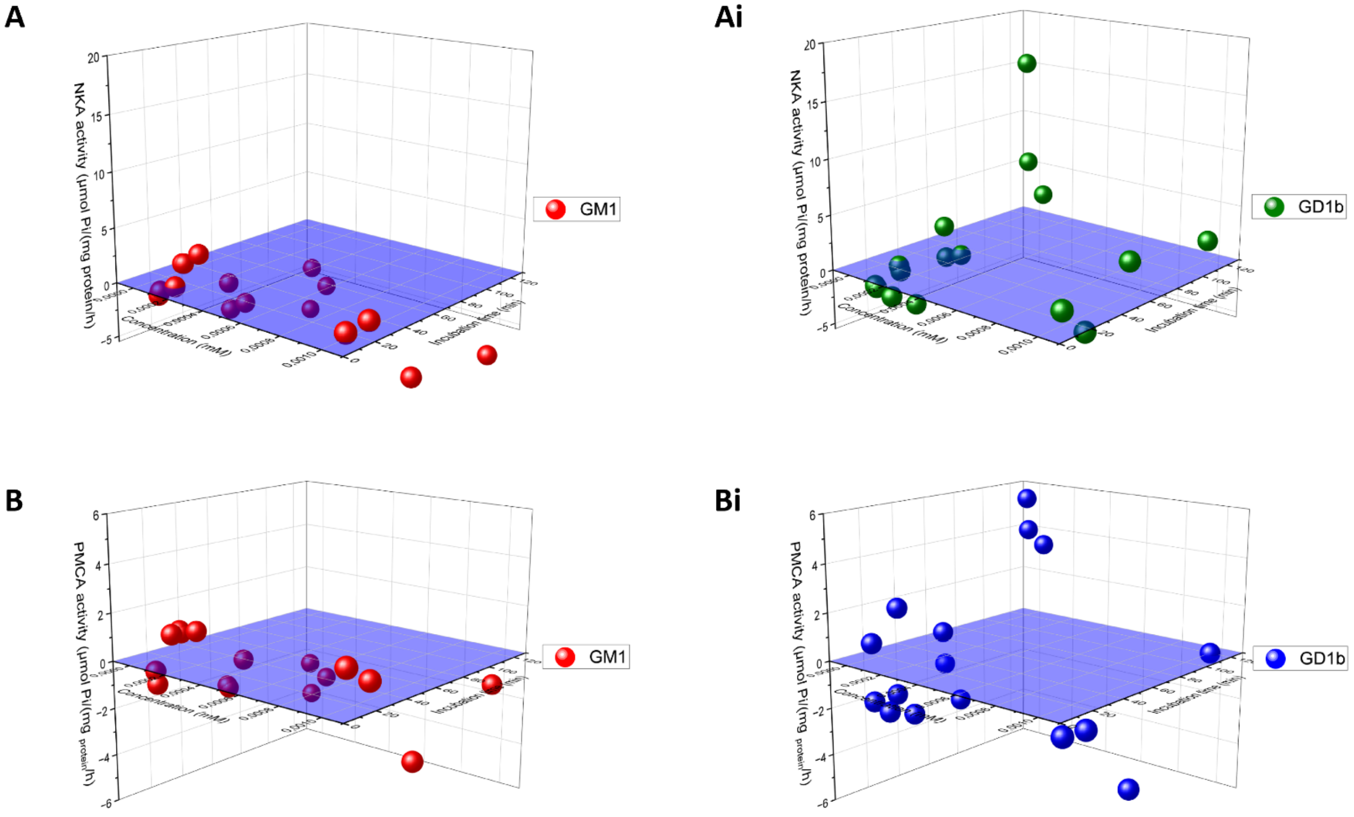


**Supplementary Figure 6.** NKA and PMCA activities in cortical homogenates of WT mice after administrations of $10^{-2}$, $10^{-3}$, $10^{-4,}$ and $10^{-6}$ mM concentrations of gangliosides GM1, GD1b and GT1b with preincubation times of 15, 30, 60 and 120 minutes. Activities are represented relative to the NKA and PMCA activities in control reactions without ganglioside administration, with 0 marking the activities in WT homogenates where no gangliosides were added. Prepared in OriginPro®2023b.

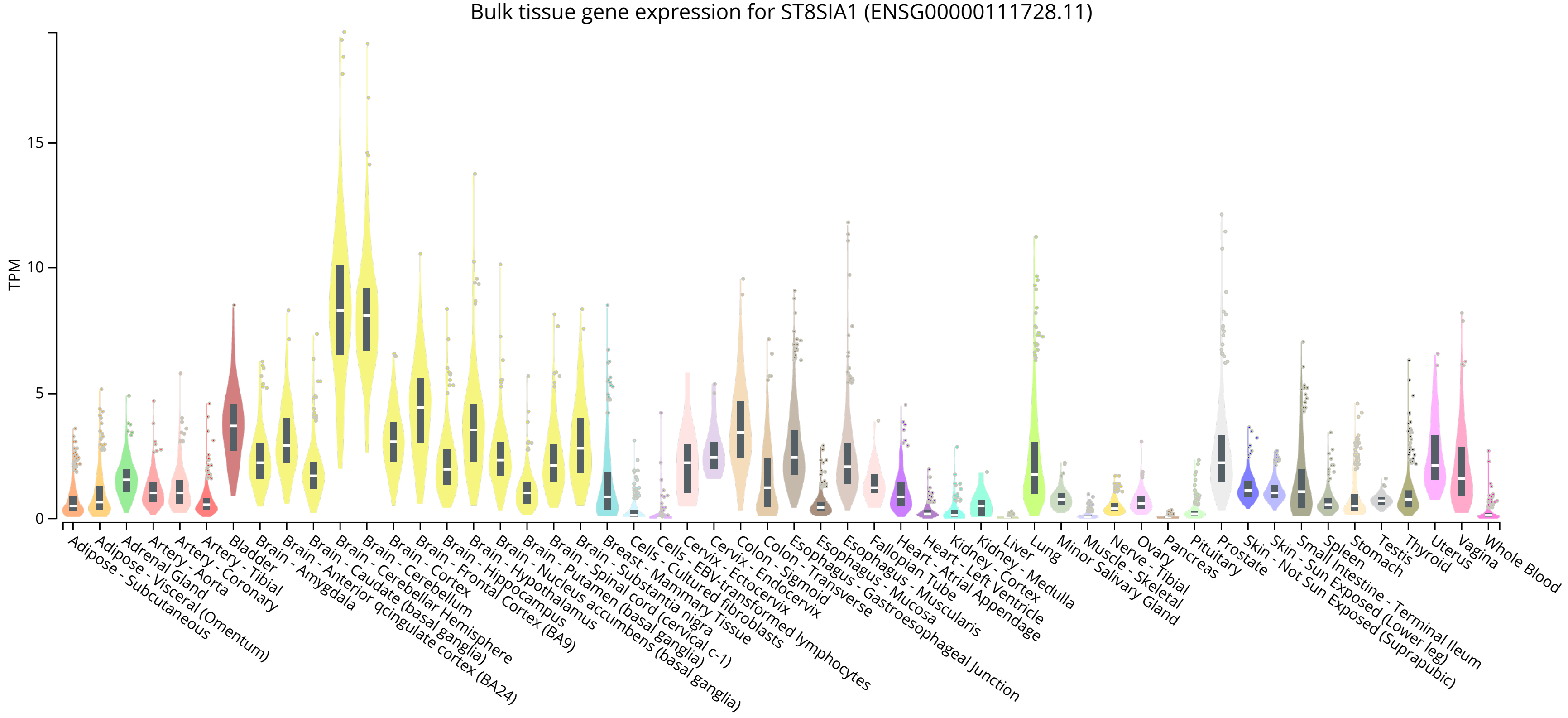


**Supplementary Figure 7.** Bulk tissue gene expression for the *ST8SIA1* gene obtained by GTEx Analysis Release V8 (dbGaP Accession phs000424.v8.p2). Expression values are shown in TPM (Transcripts Per Million), calculated from a gene model with isoforms collapsed to a single gene. Box plots are shown as median and 25th and 75th percentiles; points are displayed as outliers if they are above or below 1.5 times the interquartile range.

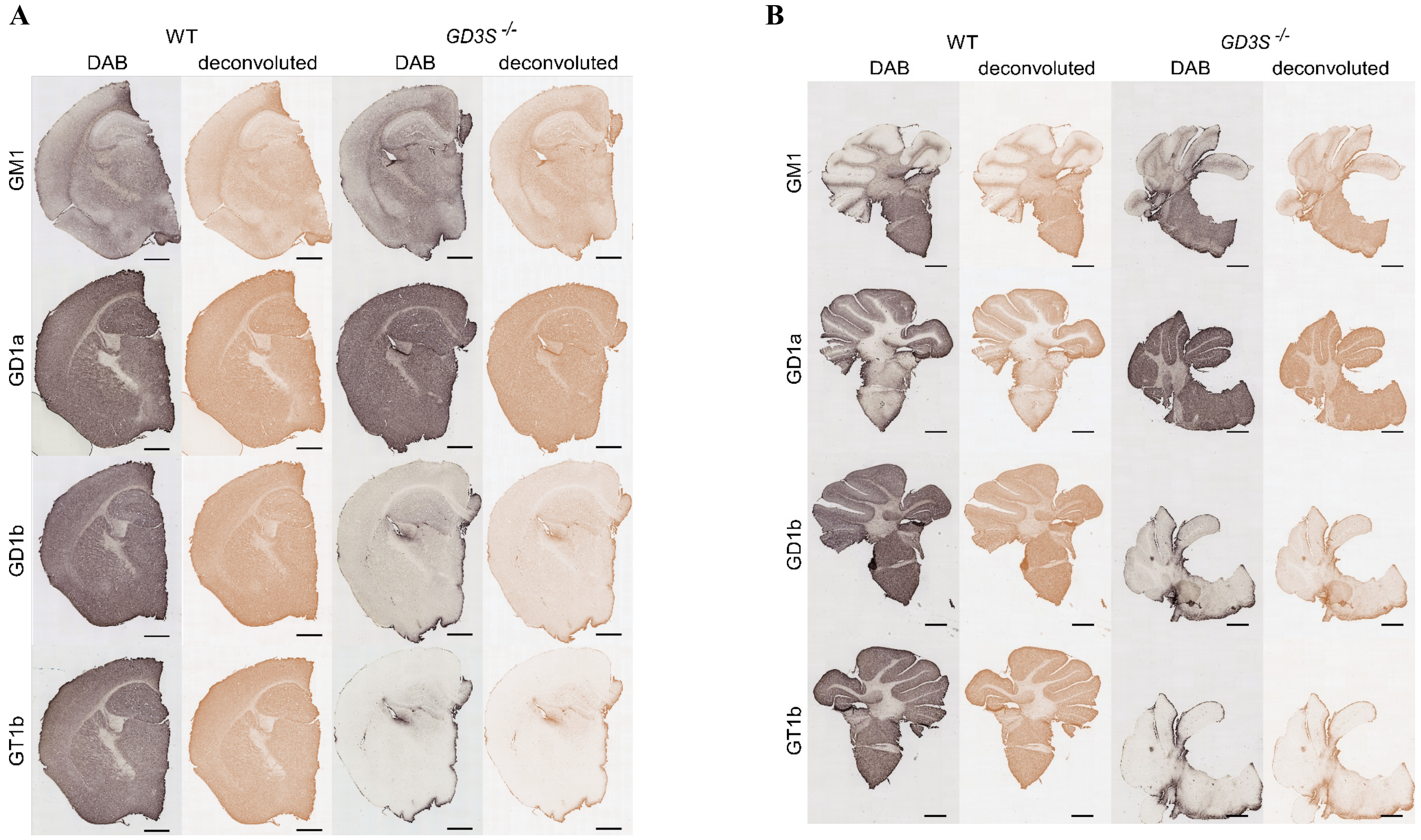


**Supplementary Figure 8.** Immunohistochemistry on four major brain gangliosides, showing no GD1b or GT1b expression in the brains of GD3S$^{-/-}$ mice. A) Cortex, scale bar = 2 mm. B) Cerebellum, scale bar = 1 mm.

**SUPPLEMENTARY TABLES**

**Supplementary Table 1.** qRT-PCR primer sequences

| Gene | Forward primer | Reverse primer | Product size (base pairs) |
|---|---|---|---|
| ***Actb*** | CATTGCTGACAGGATGCAGAA | GCTGATCCACATCTG CTGGA | 55 |
| ***Atp1a1*** | TTGAAGAGACAGCCCTTGCT | GAGGGGATACATCCTAAGGGC | 74 |
| ***Atp1a2*** | AGTCCATCGCATACACCCTG | GGAAGGGGGATGTTGGCAAT | 82 |
| ***Atp1a3*** | TCGGCTTGTTTGAGGAGACG | GAGAGGGTACATGCGAAGGG | 83 |
| ***Atp1b1*** | TTAAGAGCTGATCACAAGCACA | ACTTATTAAATGGCTAGTGGGA AAG | 50 |
| ***Atp1b2*** | TCAGCCTTTGGTGGCTGTAA | TGCGGCATTCAACATTCACC | 68 |
| ***Atp1b3*** | CATTCACAATGTGGGCCATGC | TCTGGTCTCGATATTTCGGAAC TT | 63 |
| ***Atp2b1* (Meszaros and Karin, 1993)** | TGGCAAACAACTCAGTTGCATA TAGTGG | TCCTGTTCAATTCGACTCTGCA AGCCTCG | 57 |
| ***Atp2b2*** | GCTGAACTTGGTCACACAGTC | AGTGGAGCCCATGTCTTGC | 50 |
| ***Atp2b3* (Stauffer et al., 1993)** | TCCTGTTCAATTCGACTCTGCAA GCCTCG | TCTGCTCCTGCTCAATTCGG | 68 |
| ***Atp2b4*** | CAAGCTTCGGGTACTGGCAC | TTTCACCAATGTGTGCTTGTCTG | 55 |
| ***Nptn*** | TCTCGCTCTTGCTGGTCTCT | TGGTGACAATTCTTGGTTCG | 113 |

**Supplementary Table 2. Summary of the top 25 differentially expressed genes identified from Agilent microarray analysis.** The table includes Agilent probe ID, gene symbol, full gene name, protein product, subcellular and tissue localization, and a brief description of each gene's known or proposed biological function compiled using publicly available biological databases, including NCBI Gene, UniProt, Ensembl, GeneCards, and the Human Protein Atlas.

| Agilent Probe ID | Accession ID | Gene | Full Gene Name | Protein Product | Subcellular/Tissue Localization | Function |
|---|---|---|---|---|---|---|
| A_52_P260864 | NM_009706 | *Arhgap5* | Rho GTPase activating protein 5 | p190-B RhoGAP | Cytoplasm | Regulates Rho GTPase signaling cytoskeleton remodeling |
| A_51_P446645 | XM_919321 | *Wac* | WW domain containing adaptor with coiled-coil | WAC protein | Nucleus | Transcription regulation autophagy and DNA damage response |
| A_52_P625118 | NM_007592 | *Car8* | Carbonic anhydrase 8 | Carbonic anhydrase-related protein 8 | Cytoplasm; cerebellar Purkinje cells | Regulates calcium signaling, motor coordination |
| A_51_P211526 | NM_026889 | *Ctc1* | CST telomere replication complex component 1 | CTC1 | Nucleus (telomere-associated) | Maintains telomere integrity and replication |
| A_52_P124173 | NM_172151 | *Zdhhc8* | Zinc finger DHHC-type containing 8 | Palmitoyltransferase ZDHHC8 | Golgi apparatus | Protein palmitoylation, implicated in schizophrenia |
| A_52_P174525 | BC043337 | *Epb41* | Erythrocyte membrane protein band 4.1 | Band 4.1 protein | Cytoskeleton, plasma membrane | Stabilizes membrane-cytoskeleton junctions |
| A_51_P135416 | AK165186 | *Mpped2* | Metallophosphoesterase domain-containing 2 | MPPED2 protein | Cytoplasm | Putative hydrolase, role in neuronal differentiation |
| A_51_P141012 | AK173296 | *Slitrk1* | SLIT and NTRK-like family member 1 | SLITRK1 | Plasma membrane of neurons | Promotes neurite outgrowth, linked to synapse development |

| | | | | | | |
|---|---|---|---|---|---|---|
| A_51_P156547 | NM_001079822 | *Tcf7l1* | Transcription factor 7 like 1 | TCF7L1 | Nucleus | Wnt signaling transcriptional repressor |
| A_52_P459238 | ENSMUST00000069187 | *Trim23* | Tripartite motif-containing 23 | TRIM23 (E3 ubiquitin ligase) | Cytoplasm | Ubiquitination, regulates autophagy and immune signaling |
| A_51_P211165 | BC011531 | *Rbm26* | RNA binding motif protein 26 | RNA-binding protein 26 | Nucleus (speckles) | RNA processing and splicing |
| A_51_P192089 | NM_028228 | *Pinx1* | PIN2/TERF1-interacting telomerase inhibitor 1 | PINX1 | Nucleus (telomeres) | Telomerase inhibitor, telomere length maintenance |
| A_51_P230347 | NM_177774 | *Srsf12* | Serine/arginine-rich splicing factor 12 | SRSF12 | Nucleus (spliceosomes) | Alternative splicing regulation |
| A_52_P172303 | AK031844 | *Tatdn1* | TatD DNase domain containing 1 | TatDN1 | Nucleus, cytoplasm | Putative nuclease, may regulate apoptosis/DNA repair |
| A_51_P114287 | AK077833 | *Cpsf6* | Cleavage and polyadenylation specificity factor 6 | CPSF6 | Nucleus (RNA processing foci) | 3′ end pre-mRNA cleavage and polyadenylation |
| A_51_P427080 | NM_001004364 | *Asap2* | ArfGAP with SH3 domain, ankyrin repeat and PH domain 2 | ASAP2 | Cytoplasm, plasma membrane, endosomes | Regulates membrane trafficking and cytoskeleton remodeling |
| A_51_P377237 | NM_021284 | *Kras* | Kirsten rat sarcoma viral oncogene homolog | KRAS (GTPase) | Plasma membrane (inner leaflet) | Cell signaling, growth and proliferation |
| A_51_P184728 | NM_172546 | *Cnksr* | Connector enhancer of kinase suppressor of Ras 1 | CNKSR1 | Plasma membrane & cytoplasm | Scaffold protein in Ras/MAPK signaling |

| | | | | | | |
|---|---|---|---|---|---|---|
| A_52_P338887 | NM_178761 | *Zfp672* | Zinc finger protein 672 | ZFP672 | Nucleus | Putative transcriptional regulator |
| A_52_P642005 | XM_355539 | *Camta1* | Calmodulin-binding transcription activator 1 | CAMTA1 | Nucleus | Calcium-dependent transcription factor |
| A_51_P256384 | NM_009723 | *Atp2b2* | ATPase plasma membrane $Ca^{2+}$ transporting 2 | PMCA2 | Plasma membrane (neurons, hair cells) | Calcium export pump; regulates neuronal $Ca^{2+}$ |
| A_51_P264835 | NM_010148 | *Epn2* | Epsin 2 | Epsin-2 | Cytoplasm, endocytic vesicles | Clathrin-mediated endocytosis |
| A_51_P387220 | NM_134040 | *Ddx1* | DEAD-box helicase 1 | DDX1 (RNA helicase) | Nucleus, cytoplasm | RNA processing & stress granule formation |
| A_51_P174591 | NM_012021 | *Prdx5* | Peroxiredoxin 5 | PRDX5 | Mitochondria, cytoplasm, peroxisomes | Antioxidant defense, peroxidase activity |
| A_51_P520378 | NM_007502 | *Atp1b3* | ATPase $Na^+/K^+$ transporting subunit beta 3 | $Na^+/K^+$ ATPase β3 subunit | Plasma membrane | Ion transport; stabilizes $Na^+/K^+$-ATPase complex |

### SUPPLEMENTARY MATERIALS AND METHODS

#### Immunohistochemistry

Coronal cryosections (35 µm thick) were prepared from half-brain samples for immunohistochemical analysis. Sections were initially incubated in 3% hydrogen peroxide in TBS buffer to inhibit endogenous peroxidase activity, then placed overnight at 4 °C in a blocking solution containing 1% bovine serum albumin (BSA) and 5% goat serum in TBS with gentle agitation. After blocking, sections were incubated overnight at 4 °C with primary antibodies diluted in blocking solution. These included anti-GM1 (1.1 mg/mL), anti-GD1a (1.23 mg/mL), anti-GD1b (0.36 mg/mL), and anti-GT1b (1.26 mG/mL), kindly provided by Prof. Ronald L. Schnaar, Johns Hopkins University. After washing in TBS, sections were incubated for 4 hours at 4 °C with a biotinylated goat anti-mouse secondary antibody (#115-065-166, Jackson ImmunoResearch; 1:500). Signal detection was performed using the Vectastain ABC kit (#PK-6100, Vector Laboratories) for two hours, following the manufacturer's instructions, and then washed with TBS. Visualization was performed using ROTI DAB peroxidase substrate (Carl Roth, #9202). Sections were mounted from distilled water, air-dried, and coverslipped with Biognost mounting medium (Biognost, #BM-250) for permanent preservation. Images of coronal sections were captured using a high-resolution slide scanner, the Hamamatsu NanoZoomer 2.0 RS system, with a 40× (NA 0.75) objective lens at a 455 nm/pixel resolution. For analysis, cortex images were taken at 0.5× magnification, and cerebellum images at 0.3× magnification. Images were deconvoluted using ImageJ/Fiji (NIH, USA) Image deconvolution 2 plugin with the H-DAB vector.

### SUPPLEMENTARY REFERENCES

Chen, E.Y., Tan, C.M., Kou, Y., Duan, Q., Wang, Z., Meirelles, G. V., Clark, N.R., Ma'ayan, A., 2013. Enrichr: interactive and collaborative HTML5 gene list enrichment analysis tool. BMC Bioinformatics 14. https://doi.org/10.1186/1471-2105-14-128

Meszaros, J.G., Karin, N.J., 1993. Osteoblasts express the PMCA1b isoform of the plasma membrane Ca(2+)-ATPase. J Bone Miner Res 8, 1235–1240. https://doi.org/10.1002/JBMR.5650081011

Stauffer, T.P., Hilfiker, H., Carafoli, E., Strehler, E.E., 1993. Quantitative analysis of alternative splicing options of human plasma membrane calcium pump genes. Journal of Biological Chemistry 268, 25993–26003. https://doi.org/10.1016/s0021-9258(19)74484-6
